# A Transcritical Bifurcation at Infinity Governs Plaque Stability in a Lipid-Structured Macrophage Model of Atherosclerosis

**DOI:** 10.64898/2026.01.25.701595

**Authors:** Emine Atıcı Endes, Neslihan Nesliye Pelen

## Abstract

Atherosclerotic plaques are fatty deposits in arterial walls and a major cause of heart attacks and strokes. Macrophage proliferation triggers plaque growth and instability, but the specific conditions that convert stable plaques into unstable ones remain unclear. To provide insight into the conditions for this transition, we apply bifurcation analysis to the lipid-structured atherosclerosis model proposed by Chambers et al. (Bull Math Biol 86(8):104, 2024). We demonstrate that, in the asymptotic regime where macrophage levels become large (***M* → ∞**), the model exhibits an asymptotic fast-slow structure that does not hold outside this limit. Within this asymptotic regime, we reduce the full system onto a slow invariant manifold, providing a simplified yet accurate description of the dynamics near the critical proliferation-emigration threshold. We prove, via centre manifold theory, that the positive steady state loses stability through a transcritical bifurcation at infinity at the critical proliferation-emigration threshold ***ρ***_***c***_ **= 1 + *γ***. The analysis reveals that the positive equilibrium branch approaches and exchanges stability with a boundary equilibrium at infinity, providing a rigorous dynamical explanation for the transition that was identified but left unexplored in the original study. Complementing this analytical contribution, we conduct a global sensitivity analysis using Partial Rank Correlation Coefficients (PRCC), identifying macrophage proliferation, emigration, and efferocytosis as the dominant regulators of plaque dynamics. Targeted parameter investigations near ***ρ***_***c***_ reveal a regulatory decoupling between macrophage accumulation and necrotic core growth, providing new biological insight into the behaviour of the model near the critical proliferation-emigration threshold.

## 1 Introduction

Atherosclerosis is a chronic inflammatory disease of the arteries and a leading cause of cardiovascular mortality worldwide Jebari-Benslaiman et al. (2022). The disease develops through the progressive accumulation of lipids and immune cells within the arterial wall Hansson and Libby (2006); Soleimani et al. (2021), driven by endothelial dysfunction Medina-Leyte et al. (2021), oxidative modification of low-density lipoprotein (LDL), and a self-sustaining inflammatory cycle involving monocyte recruitment Tall and Yvan-Charvet (2015), macrophage activation, and foam cell formation Chistiakov et al. (2016); Moore et al. (2013). When apoptotic macrophages are not cleared by live macrophages (as a process called efferocytosis), they undergo secondary necrosis Mehrotra and Ravichandran (2022), releasing lipids into the extracellular space and fuelling the expansion of a necrotic core. The progressive growth and destabilisation of this core is the primary driver of acute thrombotic events such as myocardial infarction and stroke Salekeen et al. (2022).

Macrophage accumulation within atherosclerotic lesions is driven by two sources: the recruitment of circulating monocytes from the bloodstream and the local proliferation of resident macrophages. While the role of recruitment is well-studied, the functional significance of local proliferation remains paradoxical and poorly understood. Although macrophage proliferation was initially observed in foam cell-rich lesions Gordon et al. (1990); Rekhter and Gordon (1995); Rosenfeld and Ross (1990), its mechanistic role remained unclear until recent work showed that proliferating macrophages are active drivers of plaque development, significantly modulating cellular dynamics and lipid accumulation Farahi et al. (2021); Lhoták et al. (2016); von Ehr et al. (2022). This body of experimental evidence presents an apparent contradiction: some studies suggest proliferation is protective by increasing the capacity for lipid clearance Yurdagul (2022); Robbins et al. (2013), while others indicate it is detrimental by expanding the pool of cells that may undergo apoptosis and contribute to the necrotic core Rosenfeld (2014); Wang et al. (2017). A critical question therefore remains unresolved: does macrophage proliferation protect plaques by enhancing necrotic core clearance, or worsen them by contributing to lipid accumulation?

Mathematical modelling provides a powerful framework for investigating the mechanisms of atherosclerosis, translating complex biological processes into predictive equations that can reveal hidden dynamics and identify critical thresholds Auley (2022); Avgerinos and Neofytou (2019); Hofmann (2024); Parton et al. (2016). These models integrate hemodynamics, such as shear stress and blood flow patterns Rostam-Alilou et al. (2022), as well as cellular dynamics, including monocyte recruitment, macrophage polarisation, and foam cell formation Chalmers et al. (2015); El Khatib et al. (2012), and molecular mechanisms, including cytokine signalling and LDL oxidation Lui and Myerscough (2021); Mukherjee et al. (2024). Within the continuum modelling tradition, lipid-structured models, which track lipid distribution across the macrophage population, offer a particularly physiologically realistic representation of plaque dynamics Chambers et al. (2023); Watson et al. (2023); Ford et al. (2019). A recent extension of this framework by Chambers et al. (2024b) incorporates macrophage proliferation via a pantograph-type non-local term in a system of partial integro-differential equations (PIDEs), with explicit focus on mid-stage atherosclerosis. A key outcome of their steady state analysis is that proliferation can reduce necrotic core lipid content by distributing the lipid load across a larger macrophage population. However, the dynamic pathways leading to these steady states, and in particular the conditions under which stability is lost, remain unexplored.

Atherosclerosis is a complex biological process involving multiple cellular and biochemical components evolving over different time-scales Thon et al. (2019). Macrophage proliferation, apoptosis, and lipid accumulation each contribute to plaque formation and stability through mechanisms that operate at distinct rates. In the Chambers et al. (2024b) model, this time-scale structure becomes analytically relevant in the asymptotic regime where the macrophage population becomes large. More precisely, as *M* → ∞, the apoptotic and necrotic variables admit a fast-relaxation structure relative to the macrophage population and live macrophage lipid load. This asymptotic separation of time-scales allows the full system to be reduced locally, near the compactified boundary corresponding to *M* → ∞, onto a slow invariant manifold Fenichel (1979); Jones (1995). The resulting reduction provides a tractable description of the dynamics near the critical proliferation-emigration threshold, rather than a global fast-slow decomposition of the model.

In this work, we exploit this asymptotic fast-slow structure together with centre manifold theory to perform a rigorous bifurcation analysis of the model proposed by Chambers et al. (2024b). We show that the positive steady state loses stability through a *transcritical bifurcation at infinity* at the critical proliferation-emigration threshold, and classify this transition using the transcritical bifurcation conditions of Kuznetsov (2004). This provides a rigorous dynamical systems interpretation of the transition identified, but left unexplored, in Chambers et al. (2024b), extending their steady state analysis to reveal the mechanism governing plaque stability.

Our paper is structured as follows:

1. Section 2 presents the dimensionless form of the lipid-structured non-local continuum model of Chambers et al. (2024b), which describes macrophage proliferation and lipid accumulation in atherosclerotic plaques.
2. Dynamical systems analysis, including equilibrium and stability properties, stability analysis through the determinant of the Jacobian matrix, and identification of a critical proliferation-emigration bifurcation, is given in Section 3.
3. Section 4 contains our main analytical contributions:
  a. Fast-slow decomposition of the system and reduction onto the slow invariant manifold (Section 4.1 and 4.2).
  b. Rigorous proof via centre manifold theory of a *transcritical bifurcation at infinity* (Section 4.3).
  c. Biological interpretation of the time-scale separation between the macrophage population and the apoptotic and necrotic core variables (Section 4.4).
4. Investigation of the influence of parameters through a global sensitivity analysis via Partial Rank Correlation Coefficients (PRCC) and targeted parameter sweeps near the critical proliferation-emigration threshold, connecting these results to biological outcomes for plaque stability, is given in Section 5.
5. Finally, we discuss all results in terms of their biological implications for plaque stability and potential therapeutic strategies, and outline directions for future research in Section 6.

## 2 Mathematical modelling

In this section, to investigate the role of macrophage proliferation in plaque evolution, we consider the lipid-structured model developed by Chambers et al. (2024b). The model consists of a non-local partial integrodifferential equation system (PIDEs) for the live and apoptotic macrophage populations, structured by intracellular lipid content, coupled to an ordinary differential equation (ODE) for the necrotic core lipid content.

Here, *M* and *P* denote the total live and apoptotic macrophage populations, respectively; *A*_*M*_ and *A*_*P*_ represent their corresponding intracellular lipid contents; and *N* denotes the necrotic core lipid content. The remaining parameters in the model are positive. Full mathematical definitions of these variables and all model parameters are given in Chambers et al. (2024b) and also Appendix A.

Applying the scalings given in Appendix A.4 to the dimensional ODE subsystem (A7), the dimensionless system governing the dynamics of the macrophage population becomes:

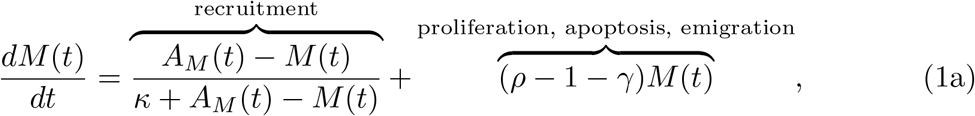

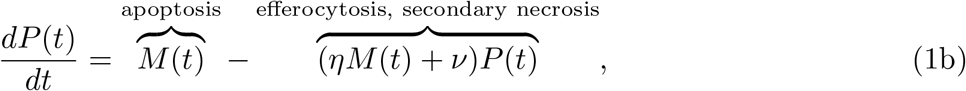

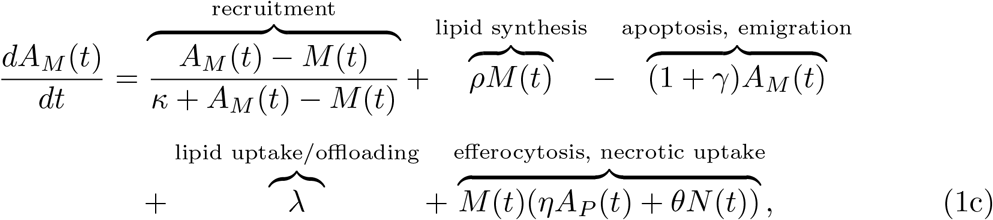

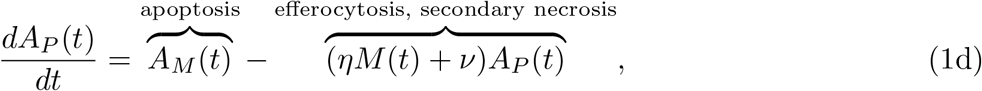

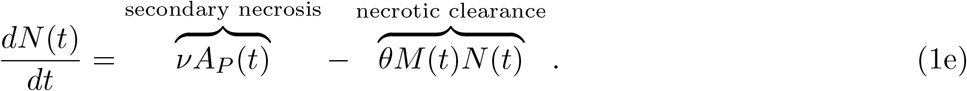

The initial conditions for the nondimensional system are given by the scaled equivalents of those in (A9):

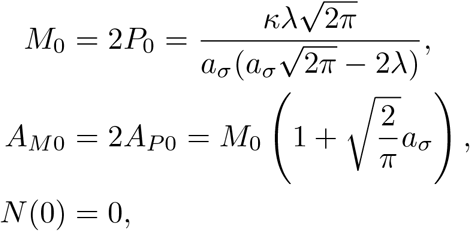

where 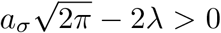 ensures biological realism. All scaled parameters are given in Section 2.1. This dimensionless system forms the basis for all subsequent analysis in this paper.

*Remark 1* It is worth noting that the dimensionless ODE system (1) cannot be obtained by directly integrating the dimensionless PDE system with respect to the lipid variable *a*. This is because Chambers et al. (2024b) employ a time-dependent normalisation, defining 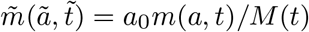 and 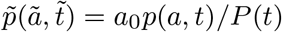, which ensures that

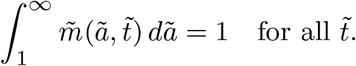

As a consequence, integrating the dimensionless PDEs over *ã* ∈ (1, ∞) yields the trivial identity 0 = 0, rather than a meaningful ODE. The dimensionless ODE system (1) is thus derived by applying the population-level scalings defined in the Appendix A.4 directly to the dimensional ODE subsystem (A7), independently of the PDE nondimensionalisation.

### 2.1 Model parameters and numerical solution

Numerical simulations and sensitivity analysis are conducted using the parameter set proposed by Chambers et al. (2024b), which supports the theoretical bifurcation analysis presented in this paper. We collect the model parameters in Table 1. In the numerical analyses presented in Section 3 and 5, we will use the parameters outlined in Table 1.

**Table 1.** Baseline parameter values used in the nondimensionalised model analysis. The parameter values are taken from Chambers et al. (2024b)

| Parameter | Biological Interpretation | Value |
| --- | --- | --- |
| $\gamma$ | Macrophage emigration rate | 0.15 |
| $\kappa$ | Half-maximal recruitment threshold | 5 |
| $\eta$ | Efferocytosis rate | 1.5 |
| $\nu$ | Secondary necrosis rate | 0.8 |
| $\theta$ | Necrotic lipid uptake rate | 0.4 |
| $\lambda$ | Net rate of lipid uptake via LDL/HDL | 0.1 |

Note: $a_\sigma$ has been chosen to be uniform. The choice of $\gamma = 0.15$ is taken to be consistent with biologically optimal regime defined by the steady state analysis of Chambers et al. \(2024b\), representing a physiologically relevant operating point near the minimum of necrotic core lipid content.

All numerical analyses were performed in Google Colab (Python 3.10) using NumPy, SciPy, scikit-learn, Matplotlib, and Seaborn. Parameter scans were conducted via Latin Hypercube Sampling, and PRCC values were computed using rank-based partial correlations with 500 bootstrap resamples. All computations and figure generations were carried out within the same Colab environment.

## 3 Dynamical systems analysis

This section presents a dynamical systems analysis of the dimensionless ODE subsystem (1). We begin by identifying the equilibrium points of the system, which represent potential stable configurations of the plaque, and establishing their stability properties through Jacobian analysis. We then perform a numerical bifurcation analysis to identify the critical proliferation-emigration threshold governing plaque dynamics. These results provide the foundation for the rigorous centre manifold analysis and bifurcation classification developed in Section 4.

### 3.1 Equilibrium analysis and stability properties

The first step in our dynamical analysis is to identify the equilibrium points of (1). To determine the equilibrium points of the system, we set the right-hand sides of equations (1a)-(1e) equal to zero and solve the resulting algebraic system. Using the steady state equations (1b)-(1e), straightforward algebraic manipulation yields the following expression for *M*\*:

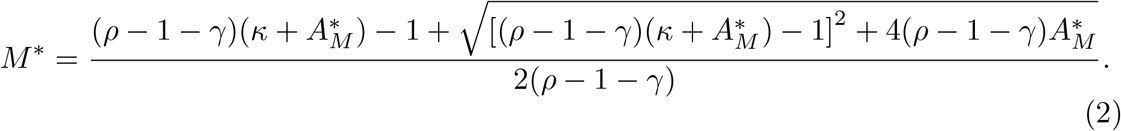

Given *M*\*, the remaining equilibrium values are determined as follows:

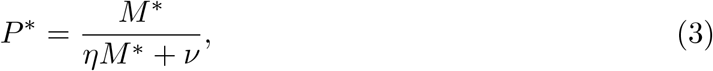

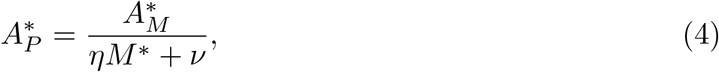

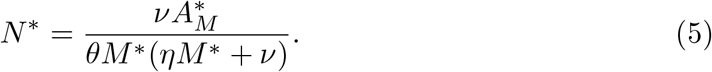

Substituting the expressions for 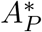 and *N*\* into (1c), the equilibrium condition for *A*_*M*_ becomes:

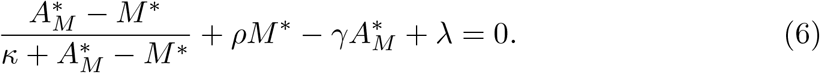

Using the equilibrium condition for *M*,

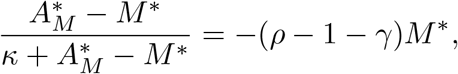

the preceding equation reduces to

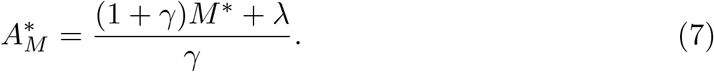

Equations (2) and (7) form a coupled algebraic system for *M*\* and 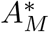. Solving these two equations determines *M*\* and 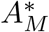, after which the remaining equilibrium values *P*\*, 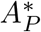, and *N*\* follow directly from (3)-(5). Furthermore, the existence of a positive equilibrium requires 1 + *γ* − *ρ >* 0, consistent with the observation of Chambers et al. (2024b).

To analyse the local stability of each equilibrium point 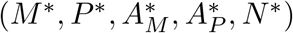, we compute the Jacobian matrix *J* of the system evaluated at the steady state. The Jacobian analysis given here serves three purposes: (i) it characterises the local stability of the equilibrium; (ii) it identifies the eigenvalue structure near the critical threshold; and (iii) it provides the foundation for the centre manifold analysis presented in Section 4.The Jacobian matrix *J* is given by:

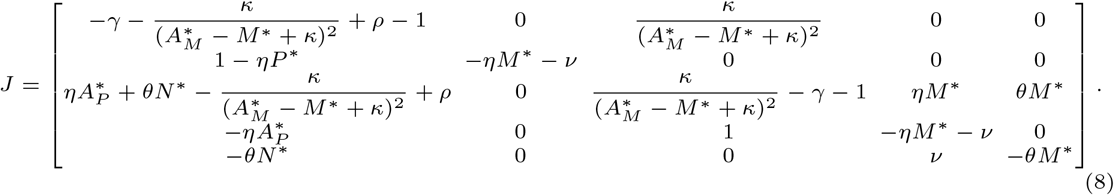

This Jacobian matrix plays a central role in the local stability analysis of the equilibrium and in identifying the eigenvalue configuration near the critical proliferation-emigration threshold. Therefore, to investigate the local stability of the equilibrium point, we compute the characteristic polynomial of the Jacobian matrix evaluated at the steady state. To write the characteristic polynomial in a simpler form, we define a new quantity:

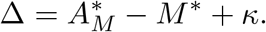

Using this definition, the characteristic polynomial *P* (*λ*) is given by

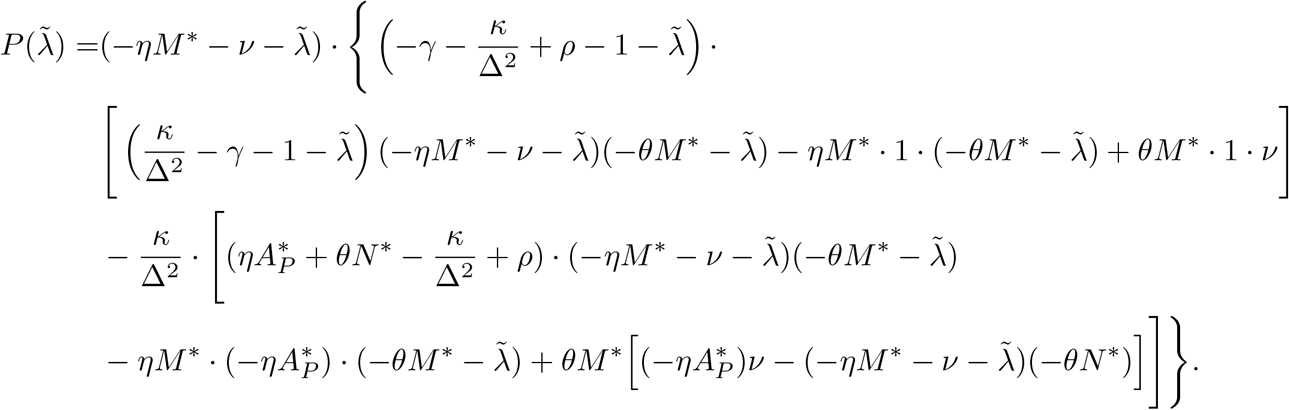

We note that the characteristic polynomial factorises as:

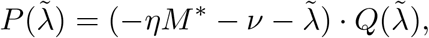

immediately identifying one eigenvalue as 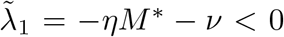. The stability of the remaining eigenvalues cannot be determined from the sign of the constant term alone for a five-dimensional system. The sign of the constant term does not provide sufficient information about the real parts of the eigenvalues. The stability question thus brings us to analysing the roots of the fourth-order factor 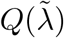, which is investigated numerically in the following subsection.

### 3.2 Bifurcation analysis

As a parameter varies, the qualitative behaviour of a system can be characterised through bifurcation analysis. The emergence, collision, or exchange of stability of equilibria at critical parameter values constitutes the core of bifurcation theory. Therefore, we next perform a bifurcation analysis to examine how these changes arise as parameters vary.

#### 3.2.1 Stability analysis through the determinant of the Jacobian matrix

Computing the determinant of the Jacobian matrix as defined in (8) allows us to analyse system stability. Figure 1 illustrates how the determinant of the Jacobian matrix changes when all parameters, as detailed in Table 1, are fixed except for *ρ*. According to the profiles in Figure 1, det(*J*) crosses zero at *ρ* = 1 + *γ*, indicating that at least one eigenvalue passes through zero. This provides numerical evidence that this value marks a genuine qualitative transition in the system, which we formally denote as

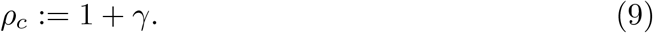

**Fig. 1.**
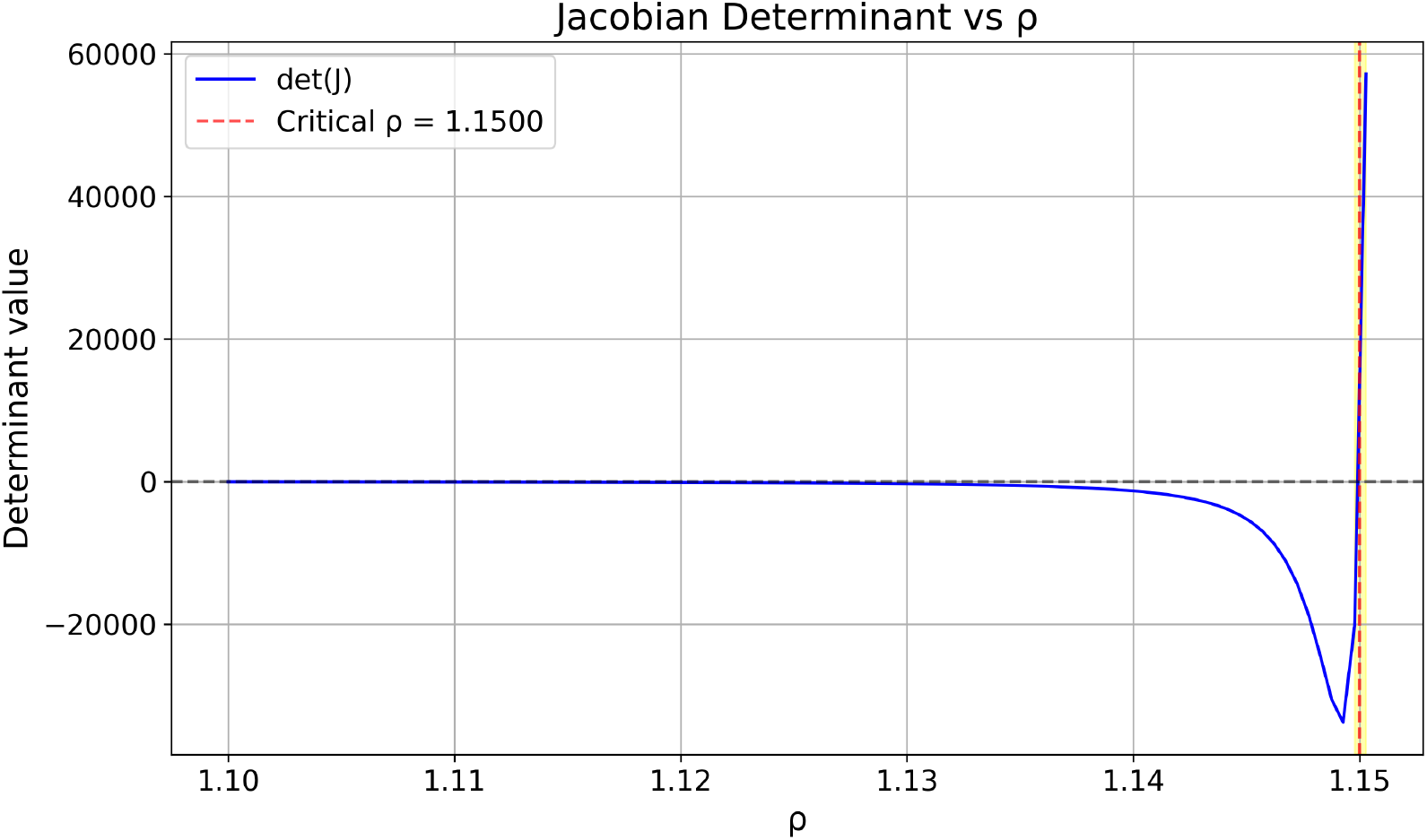
Determinant of the Jacobian matrix as a function of the macrophage proliferation parameter *ρ*. The determinant vanishes at the critical threshold *ρ* _*c*_ = 1 + *γ*, indicating the presence of a zero eigenvalue.

Consistent with the numerical simulations and the steady state analysis of Chambers et al. (2024b), the system maintains a bounded steady state for *ρ* < *ρ*_*c*_, whereas for *ρ* > *ρ*_*c*_, macrophage accumulation and lipid burden grow without bound. While Chambers et al. (2024b) imposed *ρ* < *ρ*_*c*_ as a modelling restriction without further analysis, the present work formally characterises the transition at *ρ* = *ρ*_*c*_ through determinant and eigenvalue analysis, providing the foundation for the rigorous bifurcation proof in Section 4. From a biological perspective, this transition marks a shift from controlled plaque dynamics to chronic unresolved inflammation, driven by an imbalance between macrophage proliferation *ρ* and emigration *γ*.

#### 3.2.2 Eigenvalue trajectories

In the context of bifurcation analysis, eigenvalue trajectories are essential for understanding the behaviour of dynamical systems. They illustrate how the eigenvalues of the Jacobian matrix respond to changes in specific parameters, providing insight into potential bifurcations and developments in stability. In this section, we examine Figure 2, which demonstrates the five eigenvalue trajectories of the Jacobian matrix (8) for the nondimensionalised ODE subsystem (1) as the macrophage proliferation parameter (*ρ*) varies. Each subplot corresponds to one eigenvalue, with its real (red) and imaginary (blue) parts plotted over the range of *ρ*. For clarity, a magnified view over the initial range of *ρ* is provided in Appendix C, confirming that the real parts of all eigenvalues remain negative in this region despite their apparent proximity to zero in the full-range plot.

**Fig. 2.**
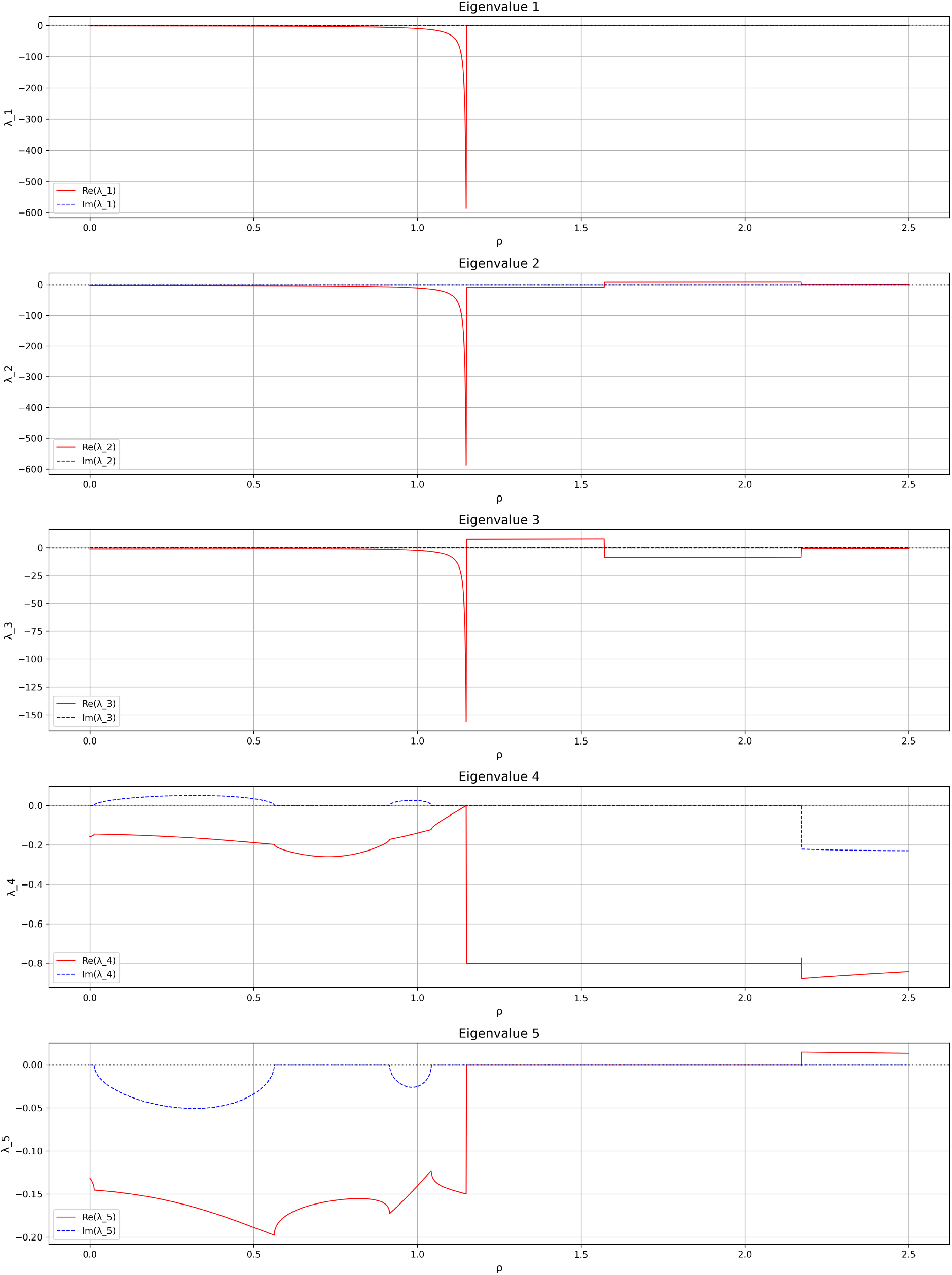
Eigenvalue trajectories near the critical point *ρ* = *ρ* _*c*_, indicating the onset of bifurcation

This analysis sheds light on how modifications in *ρ* can influence system stability and dynamic behaviour. Notably, a remarkable transition occurs around the critical point *ρ* = *ρ*_*c*_, indicating a change in the stability of the finite positive equilibrium, as some of the eigenvalues cross the imaginary axis. This crossing indicates a potential bifurcation, reflecting the central role of *ρ* in the system’s overall dynamics. Specifically:

- **Eigenvalue 1**: As the values of *ρ* vary, all these eigenvalues have large negative real parts that exhibit strongly stable directions, so there is no significant impact on bifurcation.
- **Eigenvalues 2-3**: Among these eigenvalues, Eigenvalue 3 approaches zero and its real part changes sign at *ρ* = *ρ*_*c*_, identifying the loss of hyperbolicity at the critical threshold and confirming the onset of the transcritical bifurcation at infinity established analytically. Eigenvalue 2 remains noncritical at *ρ*_*c*_, but changes sign for larger values of *ρ*, reflecting a subsequent modification of the stability spectrum beyond the bifurcation point. Beyond *ρ*_*c*_, at least one eigenvalue remains positive within the numerical spectral continuation, consistent with the onset of the post-threshold unbounded-growth regime. These post-threshold values should therefore be interpreted only as numerical indicators of the unstable regime rather than as eigenvalues outside the region of positive steady states.
- **Eigenvalues 4-5**: These eigenvalues exhibit complex behaviour with nonzero imaginary parts for small values of *ρ*, indicating damped oscillatory dynamics. They are considered together because of the related behaviour of their imaginary components. Near *ρ* = *ρ*_*c*_, their real parts remain nonpositive and do not generate the zero eigenvalue responsible for the loss of hyperbolicity. For larger values of *ρ*, the numerical spectral continuation shows a positive real part for Eigenvalue 5. Since the continuation of the nontrivial equilibrium branch lies beyond the regime of bounded macrophage accumulation for *ρ > ρ*_*c*_, this behaviour is interpreted as part of the post-threshold numerical spectrum rather than as evidence of an additional bifurcation.

Biologically, this eigenvalue analysis suggests that macrophage proliferation (*ρ*) plays a dominant role in the dynamic behaviour of the immune system. When *ρ* < *ρ*_*c*_, all eigenvalues associated with the positive finite equilibrium have negative real parts, indicating local stability. As *ρ* approaches *ρ*_*c*_, a critical eigenvalue approaches zero, signalling the loss of hyperbolicity of the finite equilibrium branch. Once the proliferation-emigration balance is disrupted, the regulatory equilibrium ceases to attract trajectories and the system evolves toward a qualitatively distinct regime, identified numerically and characterised analytically in Section 4.

#### 3.2.3 Numerical identification of the critical threshold

Having established the eigenvalue structure of the system, we now identify the critical threshold numerically. The numerical results indicate that *ρ* = *ρ*_*c*_ is the critical threshold at which the stability of the finite positive equilibrium changes, underlying a significant change in the system’s behaviour. This threshold signifies a transition in stability that is consistent with the transcritical bifurcation at infinity established analytically.

The illustration in Figure 3 is consistent with the transcritical bifurcation at infinity established analytically for system (1). This bifurcation indicates a critical transition in the system’s dynamics, as the parameter *ρ* crosses this tipping point. For *ρ* < *ρ*_*c*_, the system possesses a locally stable finite positive equilibrium. As *ρ* ↑*ρ*_*c*_, this equilibrium branch escapes to infinity. For *ρ* > *ρ*_*c*_, the nontrivial equilibrium branch continues beyond the regime of bounded macrophage accumulation, while the system evolves toward an unbounded-growth regime. Biologically, this threshold represents a balance between the growth (*ρ*) and emigration (*γ*) rates of macrophages, where the plaque stability could be affected by this disruption; for instance, increased *ρ* might simulate heightened inflammatory responses. In particular, the term (*ρ* − 1 − *γ*)*M* in equation (1a) is the key driver of this bifurcation. As *ρ* crosses the threshold *ρ*_*c*_ = 1 + *γ*, this coefficient changes sign, removing the effective damping on the macrophage population. Consequently, *M* grows rapidly, which in turn drives the increase in *A*_*M*_. The remaining variables *P, A*_*P*_, and *N* are subsequently affected through their coupling with these dominant components. Consequently, this simulation shows how model behaviour is altered under the critical parameter regime, offering significant insights into the dynamics of atherosclerosis.

**Fig. 3.**
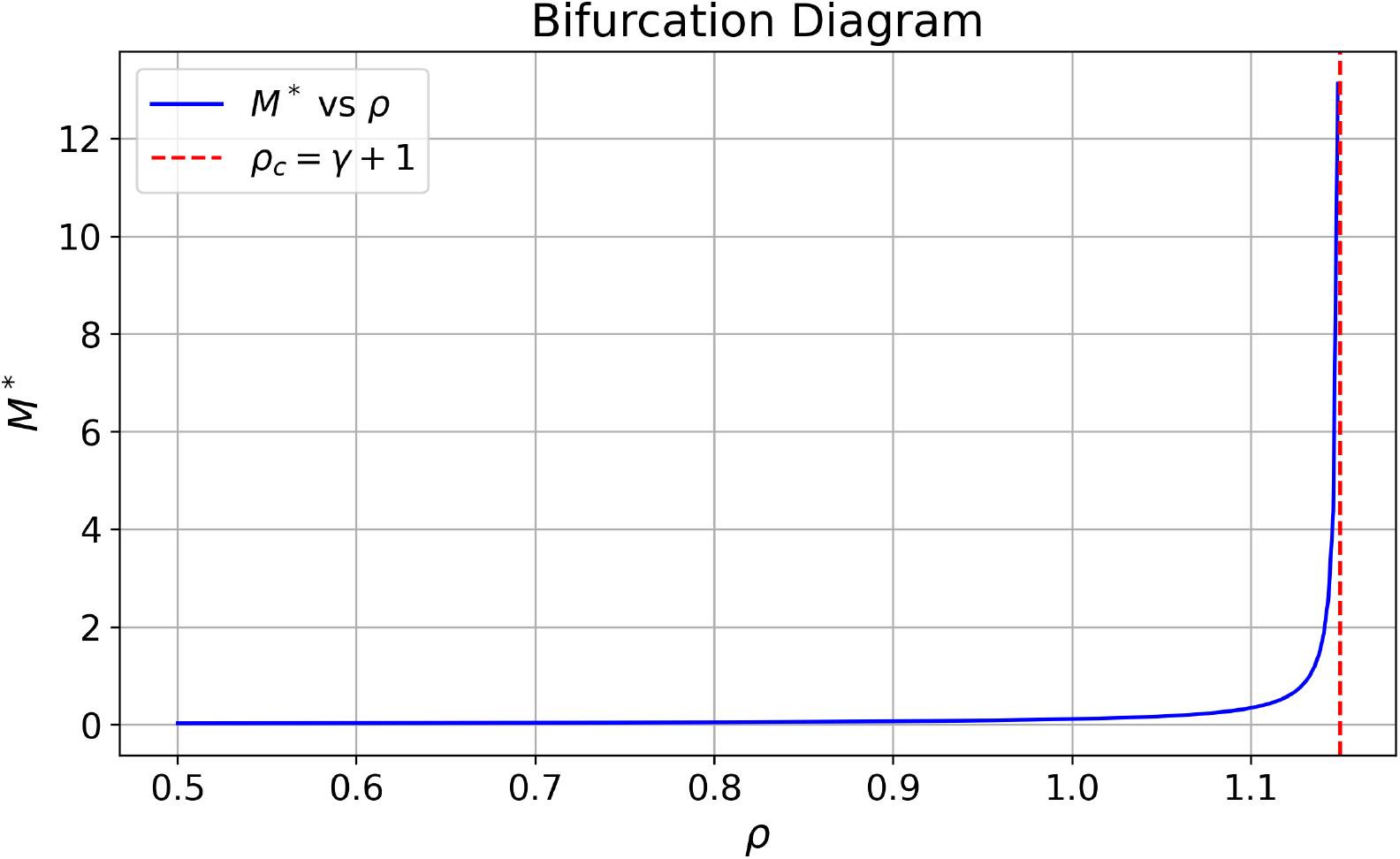
Transcritical bifurcation at infinity occurring at the threshold *ρ* = *ρ*_*c*_. As *ρ* ↑ *ρ*_*c*_, the finite positive equilibrium satisfies *M*\* → ∞, approaching the equilibrium at infinity where the transcritical bifurcation occurs.

## 4 Time-scale analysis and centre manifold theory

The numerical results of Section 3 confirm that the system undergoes a qualitative transition at *ρ* = *ρ*_*c*_, associated with the loss of stability of the finite positive equilibrium and the emergence of an unbounded growth regime. However, due to the system’s high dimensionality, a direct analytical characterisation of this transition is difficult. To overcome this difficulty, we exploit the asymptotic separation of time-scales that emerges in the large-macrophage regime (*u* → 0^+^, equivalently *M* → ∞). Within this asymptotic regime, the dynamics can be reduced to a lower-dimensional slow subsystem, where the bifurcation structure can be analysed rigorously. We show that this transition corresponds to a *transcritical bifurcation at infinity* at *ρ* = *ρ*_*c*_, and provide a rigorous proof via centre manifold theory. We emphasise at the outset that this fast-slow separation is a strictly local, asymptotic property that emerges only in the singular limit *u* → 0^+^ (equivalently *M* → ∞). It does not describe the dynamics of the bounded trajectories shown in Figures 5 and 8, which lie in the regime *ρ* < *ρ*_*c*_ away from this limit. Outside the asymptotic regime, the apoptotic and necrotic variables are not naturally fast unless the corresponding rate parameters *η, θ* are taken large.

### 4.1 Fast-slow decomposition of the system

To identify the asymptotic time-scale structure of the model near the unbounded-macrophage regime, we introduce the compactifying variables

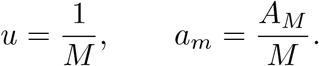

Thus, the limit *u* → 0^+^ corresponds to *M* → +∞. Using *M* = 1*/u* and *A*_*M*_ = *a*_*m*_*/u*, the equations governing the transformed variables become

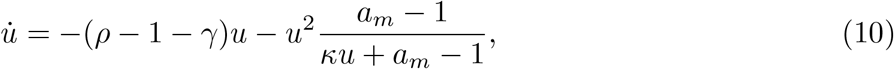

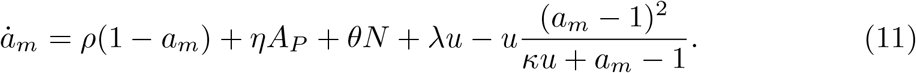

The remaining variables satisfy

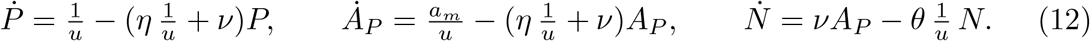

The factors 1*/u* in these equations indicate that (*P, A*_*P*_, *N*) relax on a faster time-scale than (*u, a*_*m*_) as *u* → 0^+^. To make this separation explicit, we introduce the fast time variable *τ* by

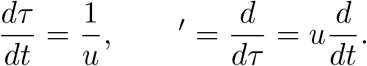

In the fast time-scale, system (12) becomes

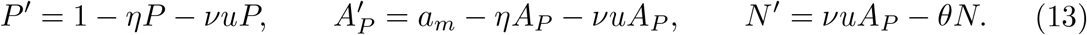

The transformed slow variables satisfy

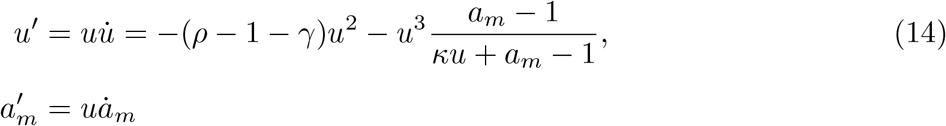

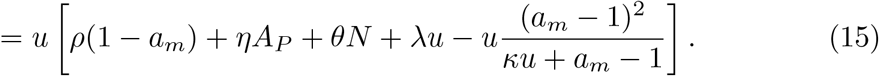

Consequently, *u*′ = 0, 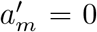 at *u* = 0. Thus, in the layer problem obtained by setting *u* = 0, the variables *u* and *a*_*m*_ are frozen, whereas (*P, A*_*P*_, *N*) retain nontrivial fast dynamics. The corresponding layer problem is

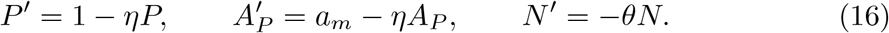

This establishes the local asymptotic fast-slow structure of the system near *u* = 0^+^. We emphasise that this decomposition is valid only in the large-macrophage limit *M* → +∞, and does not constitute a global time-scale separation for bounded values of *M*.

### 4.2 Reduction onto the slow invariant manifold

The fast-slow formulation established in Section 4.1 enables a rigorous reduction of the full five-dimensional system near the singular limit *u* → 0^+^. In the fast time-scale, the layer problem is given by

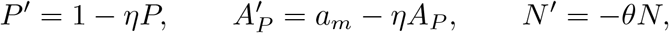

where the slow variables *u* and *a*_*m*_ are treated as frozen parameters. The layer problem possesses the unique equilibrium

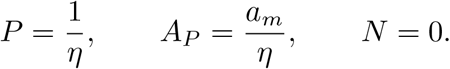

Consequently, the associated critical manifold is

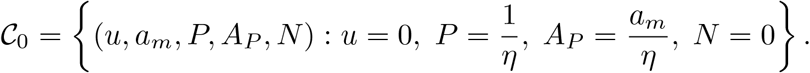

#### Lemma 1

(Normal hyperbolicity of the fast block near *u* = 0^+^) *Let u* ∈ [0, *u*_0_] *and let am belong to a compact interval I* ⊂ ℝ. *Assume that η, ν, θ* > 0. *Consider the fast subsystem in* (13). *For each fixed* (*u, am*), *this subsystem has the unique equilibrium*

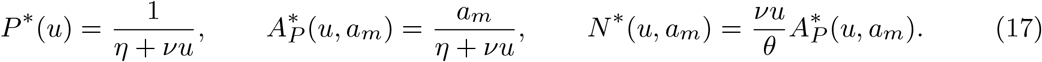

*The Jacobian of the fast block at* 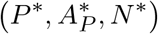 *is*

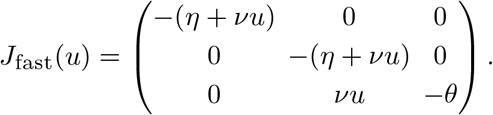

*Hence, its eigenvalues are*

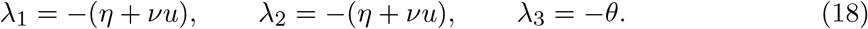

*In particular, there exists a constant c* > 0, *independent of u* ∈ [0, *u*_0_], *such that*

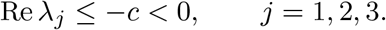

*At u* = 0, *the family of fast equilibria reduces to the critical manifold*

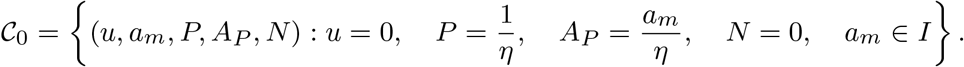

*Therefore, C*_0_ *is normally hyperbolic and attracting*.

*Proof* For fixed (*u, am*), the equilibrium is obtained by solving

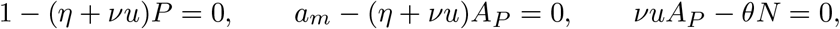

which gives (17).

The eigenvalues of the Jacobian matrix are its diagonal entries:

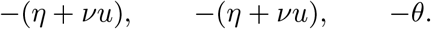

Since *η, θ* > 0 and *u* ≥ 0, these eigenvalues are uniformly bounded away from the imaginary axis. Indeed, one may take

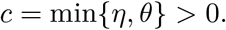

Thus all fast directions are uniformly attracting.

Taking the singular limit *u* = 0 in (17) gives

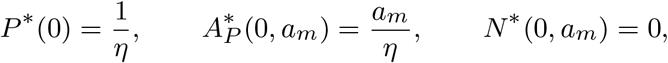

which defines *C*_0_. Hence *C*_0_ is normally hyperbolic and attracting.

By Fenichel’s invariant manifold theorem (Theorem 4 in Appendix B), every compact portion of the normally hyperbolic critical manifold *C*_0_ persists, for sufficiently small *u* > 0, as a locally invariant attracting slow manifold *C*_*u*_, which is *C*^*r*^-close to *C*_0_.

On this perturbed slow manifold, the fast variables admit smooth expansions in terms of the slow variables. Solving the fast equilibrium equations for *u* > 0 gives

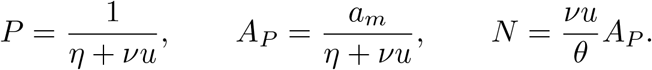

Since the invariant slow manifold is *C*^*r*^-close to the critical manifold *C*_0_, its leading-order approximation is obtained from the equilibrium of the frozen fast subsystem. Consequently,

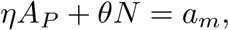

and therefore

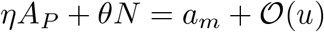

on the invariant slow manifold.

Substituting these quasi-steady expressions into the remaining equations yields an effective two-dimensional reduced system

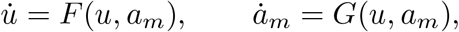

which governs the slow dynamics of the original system in a neighbourhood of *u* = 0^+^.

The occurrence of a *transcritical bifurcation at infinity* was numerically identified in Section 3.2.3; see Figure 3. The reduced slow system obtained above provides the appropriate framework for analysing this transition rigorously. In the following subsection, we study the reduced dynamics near the singular point

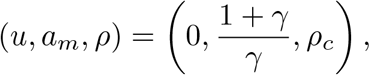

and prove that the observed transition is a transcritical bifurcation at infinity.

As established in Section 3.2.1, for all *ρ* < *ρ*_*c*_ := 1 + *γ* the system admits a unique positive equilibrium 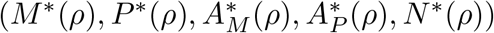. Moreover, by using Equations from (2) to (7), one can also show that there is a unique equilibrium branch for *ρ* < *ρ*_*c*_,

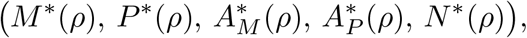

such that *M*\*(*ρ*) → +∞, 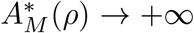 and 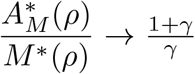 as *ρ* ↑*ρ*. Now, we take these results as a starting point for our analytical bifurcation analysis. Here *ρ*_*c*_ gives the critical value of *ρ*. To formalise the analysis, we introduce the following assumptions.

Assumptions

**(H1)** Parameters satisfy *κ, η, ν, θ, λ, γ* > 0. Write *ρ* = *ρ*_*c*_ + *µ* with *ρ*_*c*_ := 1 + *γ* and |*µ*| < *µ*_0_ for some *µ*_0_ > 0.

**(H2)** With the given initial/boundary data, the positive cone is forward invariant and the vector field is at least *C*^2^ on the relevant domain.

The significance of Theorem 2 is that it provides a rigorous mathematical explanation for the threshold previously observed only numerically. Rather than simply identifying the loss of stability of the finite positive equilibrium, it establishes that this transition occurs through an exchange of stability with the boundary equilibrium at infinity, thereby characterising the underlying mechanism as a transcritical bifurcation at infinity.

#### Theorem 2

(Transcritical bifurcation at infinity) *Assume* ***(H1)*** *and* ***(H2)***. *Introduce u* := 1*/M, a* _*m*_ := *A*_*M*_ */M, and µ* := *ρ* − *ρ* _*c*_, *so u* → 0 *corresponds to M* → ∞. *Then the system undergoes transcritical bifurcation at ρ* = *ρ*_*c*_ *at infinity*.

The proof of Theorem 2 is given in Section 4.3.

#### Remark 2

The *transcritical bifurcation at infinity* identifies the critical proliferation-emigration threshold at which the finite positive equilibrium loses stability through an exchange of stability with the equilibrium at infinity. Biologically, this transition marks the point beyond which macrophage proliferation dominates the regulatory mechanisms that normally maintain plaque homeostasis. Consequently, the model predicts the onset of an unbounded inflammatory regime associated with progressive plaque growth and loss of plaque stability.

### 4.3 Centre manifold analysis and proof of the transcritical bifurcation at infinity

#### Proposition 3

*[`Fast-slow decomposition and slow manifold near` u* = 0^+^*] Let u* := 1*/M and a*_*m*_ := *A*_*M*_ */M. For* (*u, a*_*m*_) *in a neighborhood of* 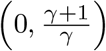 *and parameters as in* (H1)-(H2), *consider the fast block* (*P, A*_*P*_, *N*) *in subsystem* (13). *Then there exist u*_0_ > 0 *and a neighborhood U of* 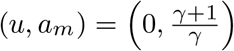 *such that for all* 0 < *u* ≤ u0 *and* (*u, am*) ∈ *U:*

1. *The fast block has a unique equilibrium given as in* (17) *and its Jacobian has a spectrum as in* (18), *hence is uniformly normally hyperbolic and attracting with a spectral gap bounded below by some constant c* > 0.
2. *The critical manifold*

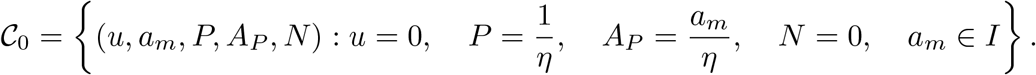

*persists to an attracting C*^*k*^ *such that k* ≥ 2 s*low manifold C*_*u*_ *for u* ∈ (0, *u*_0_], *with the expansion*

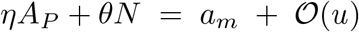

*uniformly on compact subsets of U*.

*Proof* Part (1) follows from Lemma 1. Solving the steady state equations yields the equilibrium 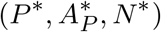, and the Jacobian, which is lower triangular, has eigenvalues given by (18). Since

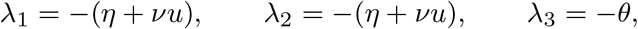

there exists a constant

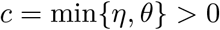

such that

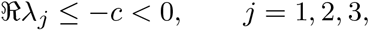

for all *u* ∈ [0, *u*_0_]. Hence, the fast eigenvalues are uniformly bounded away from the imaginary axis, which establishes uniform normal hyperbolicity and attraction.

For Part (2), the vector field is *C*^*k*^ by (H2) and the normal hyperbolicity in Part (1) is uniform for *u* ∈ (0, *u*_0_] and (*u, a*_*m*_) near 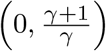. Hence, Theorem 4 in Appendix B yields an attracting *C*^*k*^slow manifold *C*_*u*_ that is *O*(*u*)-close (in *C*^0^) to *C*_0_. Evaluating along *C*_*u*_ gives *ηA*_*P*_ + *θN* = *a*_*m*_ + *O*(*u*) uniformly on compact subsets.

Now by using the above Lemma 1 and Proposition 3 we will prove **Theorem 2** as follows:

*Proof* To analyse the local dynamics near the critical point, we introduce a blow-up coordinate near 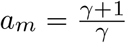 by setting

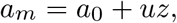

where

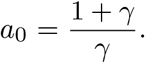

Using (10), the *u*-equation takes the form

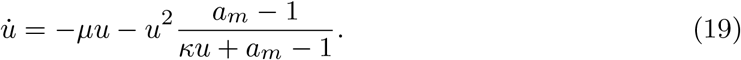

In the original time-scale, the fast subsystem is given by

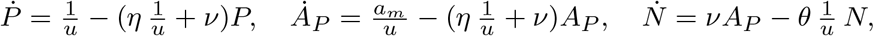

using the quasi-steady relations derived earlier,

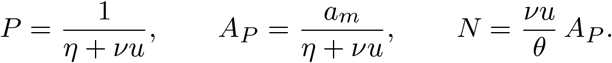

The quasi-steady relations above satisfy the identity

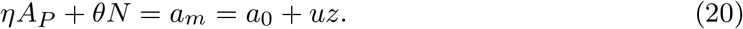

At *u* = 0, these relations reduce to the critical manifold

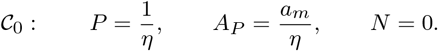

By Lemma 1, the fast block is uniformly normally hyperbolic for sufficiently small *u* > 0. Hence Fenichel theory Fenichel (1979); Jones (1995) guarantees the existence of a locally invariant attracting slow manifold *C*_*u*_ satisfying

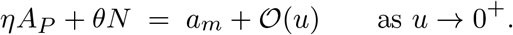

Since

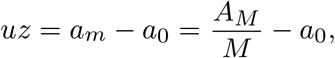

we introduce the translated variable *v* := *am* − *a*_0_ = *uz*. Proposition 3 ensures that the attracting slow manifold obtained after eliminating the fast variables is of class *C*^*r*^, with *r* ≥ 2. Since the coordinate transformation

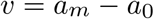

is a *C*^∞^ translation, the resulting reduced vector field in the variables (*u, v, µ*) is also of class C^*r*^. We augment the reduced system by treating the bifurcation parameter as a dynamical variable,

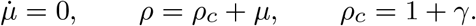

Since

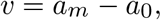

we have 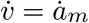. On the attracting slow manifold, we have

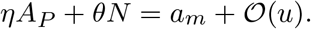

Substituting *a*_*m*_ = *a*_0_ + *v* into (11) gives

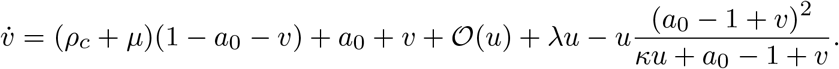

To compute the linearisation explicitly, first recall that

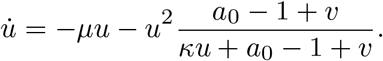

Since both terms on the right-hand side are at least quadratic in (*u, v, µ*), the *u*-equation has no linear terms at (*u, v, µ*) = (0, 0, 0). Hence 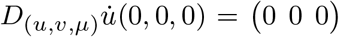. Next, expanding the terms in the *v*-equation gives

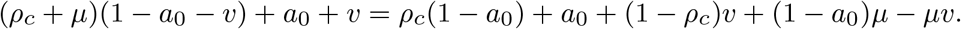

Using

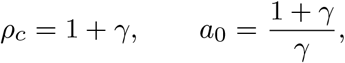

we obtain

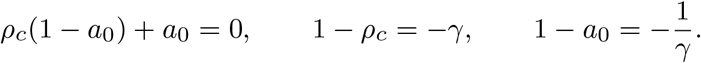

Therefore,

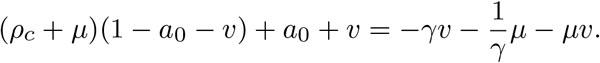

Moreover,

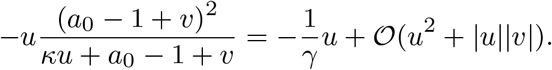

Since the reduced vector field is of class *C*^*r*^, the remainder *O*(*u*) arising from the slow-manifold reduction may be written locally as

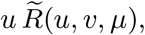

where 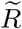 is continuous, and in fact *C*^*r*−1^. Consequently, the linearised *v*-equation has the form

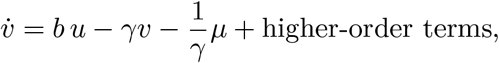

where

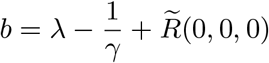

collects all linear contributions proportional to *u*. Together with 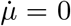, the Jacobian of the extended system at the origin is therefore

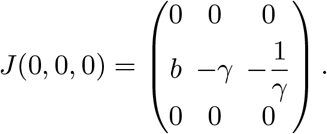

The eigenvalues of this matrix are:

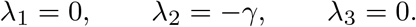

Since *γ* > 0, the eigenvalue −*γ* is strictly negative, whereas the remaining two eigenvalues are zero. Thus the system has a two-dimensional centre subspace and a one-dimensional stable subspace. The stable eigenspace is the *v*-axis, and the centre subspace is transverse to it. Therefore, the centre manifold theorem guarantees the existence of a locally invariant centre manifold that can be represented as

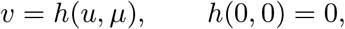

with *h* ∈ *C*^*r*^. Since *h* ∈ *C*^*r*^ with *r* ≥ 1, the first-order Taylor expansion of *h* at (0, 0) implies

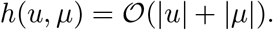

Define

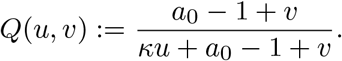

Since

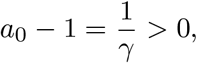

*Q* is smooth near (0, 0), its first-order Taylor expansion about (0, 0) gives

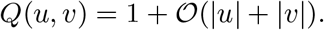

Since

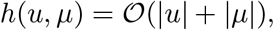

it follows that

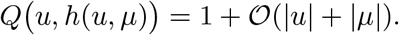

Substituting *a*_*m*_ = *a*_0_ + *h*(*u, µ*) into (19) gives

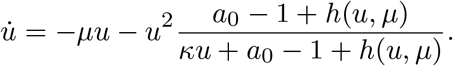

By the definition of *Q*, 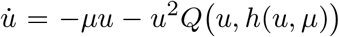. Since *Q u, h*(*u, µ*) = 1 + *O*(|*u*| + |*µ*|), we obtain

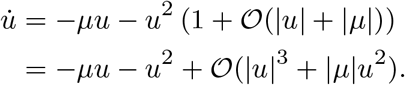

Hence, the reduced equation on the centre manifold becomes

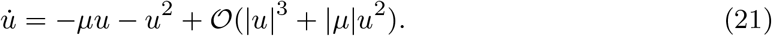

Let *F* (*u, µ*) = −*µu* − *u*^2^ + *O*(|*u*|^3^ + |*µ*|*u*^2^). Then

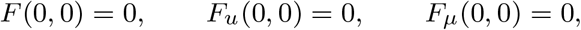

while *F*_*uµ*_(0, 0) = −1≠ 0, *F*_*uu*_(0, 0) = −2 ≠ 0. These are the standard nondegeneracy conditions for a transcritical bifurcation. Since every term of *F* contains the factor *u*, we write

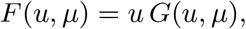

where *G*(*u, µ*) = −*µ*−*u*+*O*(*u*^2^+|*µ*||*u*|). The nontrivial equilibria therefore satisfy *G*(*u, µ*) = 0. Since the derivative of *G*(*u, µ*) with respect to *u* at (*u, µ*) = (0, 0) is −1 ≠ 0 and *G*(0, 0) = 0 the implicit function theorem yields a unique local nontrivial equilibrium branch

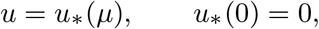

with *u*\*(µ) = −*µ* + *O*(*µ*^2^). Along the trivial branch,

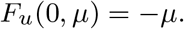

Along the nontrivial branch, using *u*\*(µ) = −*µ* + *O*(*µ*^2^), we obtain

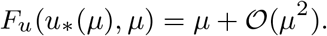

Hence the two equilibrium branches exchange stability as *µ* passes through zero. Consequently, the reduced system undergoes a transcritical bifurcation at

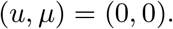

In the original variables, this corresponds to a transcritical bifurcation at infinity at

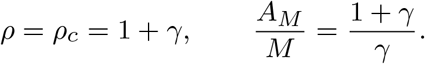

### 4.4 Biological significance of the time-scale separation

Based on the asymptotic large-macrophage regime analysed here, the fast-slow structure indicates that the apoptotic and necrotic components, represented by *P, A*_*P*_, and *N*, adjust rapidly to changes in the system and remain close to their quasi-steady states. In contrast, the macrophage population *M* and live macrophage lipid content *A*_*M*_ evolve on a slower time-scale and therefore determine the overall progression of the system. Considering this asymptotic regime, this implies that plaque progression is primarily governed by macrophage proliferation and lipid accumulation dynamics, while apoptotic and necrotic processes rapidly relax to their quasi-steady states. This suggests that interventions targeting macrophage proliferation and emigration rates may play a key role in influencing long-term disease progression in this regime.

Moreover, within the asymptotic regime considered here, the presence of a *transcritical bifurcation at infinity* reveals an underlying threshold mechanism: gradual changes in macrophage proliferation and emigration can induce a precise qualitative transfer of stability associated with plaque destabilisation. This provides a biological interpretation in which macrophage accumulation and lipid processing play a central role in driving plaque progression, while the existence of a critical threshold *ρ*_*c*_ = 1 + *γ* offers a mathematical explanation for the qualitative change in plaque dynamics observed in advanced atherosclerosis, and identifies the conditions under which this transition may be reversed.

## 5 Parameter analysis and biological implications

This section investigates the biological consequences of the transcritical bifurcation at infinity established in Section 4. We begin with a global sensitivity analysis to identify the dominant parameters governing system behaviour. Building on these insights, we then conduct targeted parameter investigations characterising how these dominant parameters influence the transition from stable plaque dynamics to unbounded macrophage accumulation near the critical threshold *ρ* = *ρ*_*c*_.

### 5.1 Parameter sensitivity-PRCC method

In this section, we provide a systematic framework for identifying the key biological mechanisms governing plaque dynamics and for assessing the relative importance of macrophage proliferation, emigration, and efferocytosis in shaping system behaviour. To do that, we perform a global sensitivity analysis (GSA) to quantify the influence of model parameters on system outputs. Unlike local sensitivity approaches, which assess parameter effects around a fixed reference point, global sensitivity analysis evaluates the impact of parameter variations across their full physiological ranges Li et al. (2023); Qin et al. (2023). This is particularly important for capturing nonlinear interactions inherent in biological systems Kent et al. (2013); Marino et al. (2008).

We specifically use the Partial Rank Correlation Coefficient (PRCC) method Gasior (2025); Sorokin and Goryanin (2023). PRCC is a powerful GSA approach that quantifies the monotonic relationship between each input parameter and each model output, while controlling for the influence of all other parameters, making it a robust tool for identifying the dominant drivers of system behaviour. The parameters are sampled over biologically relevant ranges, such that *κ* in [0.5, 10], *ρ* in [0.8, 1.5], *γ* in [0.05, 0.3], *λ* in [0.05, 0.3], *η* in [0.05, 3], *ν* in [0.05, 1], and *θ* in [0.05, 1], using Latin Hypercube Sampling (LHS), generating 500 parameter sets. For each sampled set, the model is simulated up to *t* = 50 using an explicit Euler scheme with time step Δt = 0.05, and the terminal values of *M*, *P*, *A*_*M*_, *A*_*P*_, and *N* are recorded. The results are summarised in Figure 4, where rows represent the input parameters and columns correspond to the model outputs *M, P, A*_*M*_, *A*_*P*_, and *N*.

**Fig. 4.**
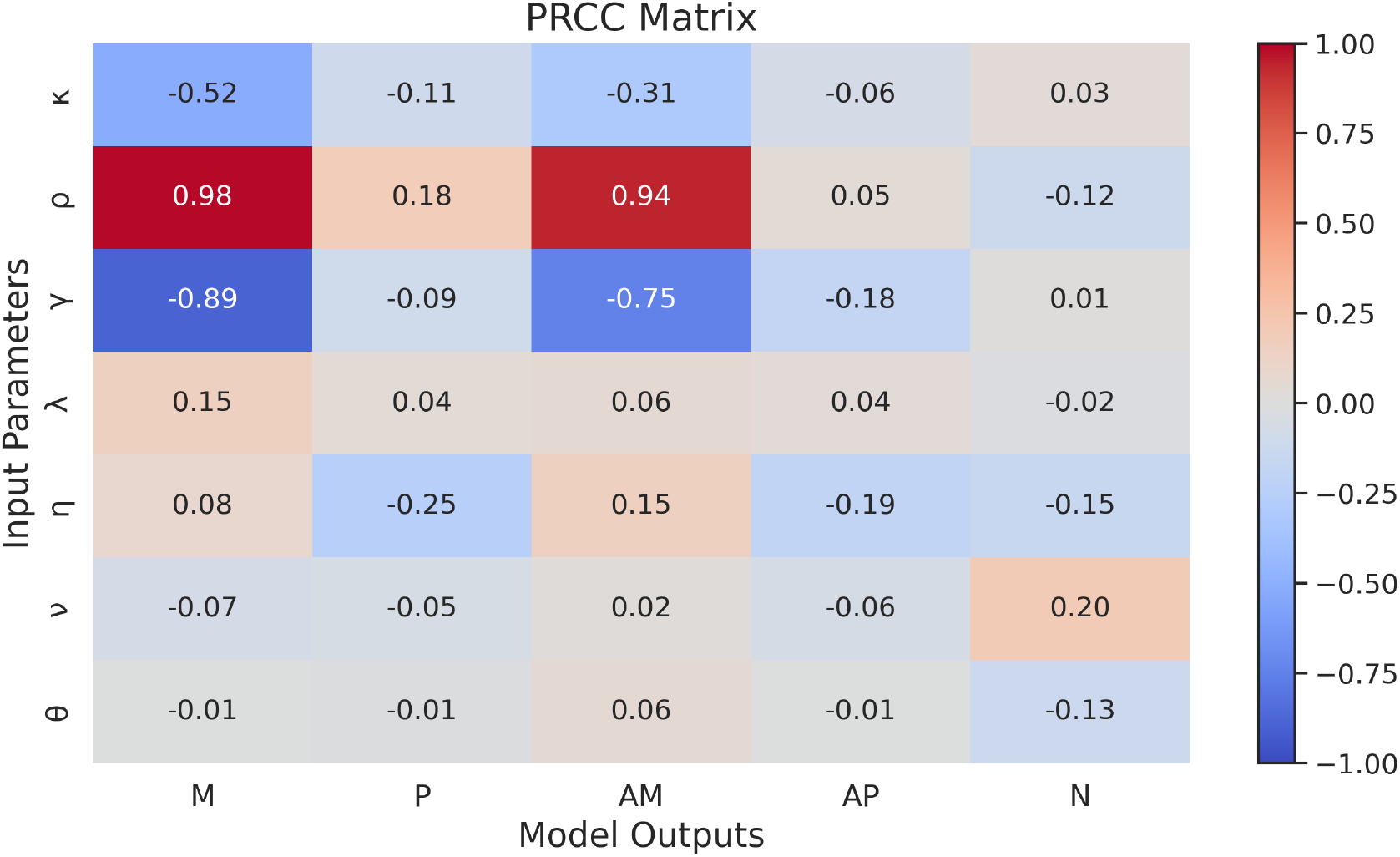
Partial Rank Correlation Coefficient (PRCC) heatmap illustrating the sensitivity of model outputs (*M, P, A*_*M*_, *A*_*P*_, *N*) to variations in the most influential input parameters (*ρ, γ, η*). Red shades represent strong positive correlations, blue shades represent strong negative correlations, and near-zero values indicate weak, meaning negligible influence

The magnitude and sign of the PRCC values indicate the strength and direction of the relationship between parameters and model outputs. As long as the PRCC values are close to +1 or −1, one can say that there is a powerful relation between the parameter and the considered output. However, if the PRCC value is near zero, this shows little influence of the parameter on the output. Additionally, the sign of the PRCC indicates the direction of the parameter’s effect. If the sign is positive, as the parameter rises, the considered output also increases. However, when the sign is negative, even as the parameter increases, the output value decreases.

From the heatmap, several observations can be made:

- The macrophage proliferation rate *ρ* exhibits near-unity positive correlations with *M* and *A*_*M*_ (approximately 0.98 and 0.94), indicating that the system operates in a proliferation-dominated regime in which macrophage growth governs both population size and live macrophage lipid content.
- The emigration rate *γ* exhibits strong negative correlations with *M* and *A*_*M*_, indicating that macrophage removal acts as a dominant suppressive mechanism that counterbalances accumulation and lipid loading. This highlights the role of emigration as a key regulatory pathway limiting macrophage-driven plaque growth.
- The efferocytosis rate *η* exhibits an asymmetric influence across the model outputs. Its most pronounced negative correlations occur with the apoptotic compartments *P* and *A*_*P*_, while its effects on the remaining variables are comparatively weaker, including a modest positive correlation with the live macrophage lipid content *A*_*M*_. This pattern is consistent with the primary role of efferocytosis in regulating apoptotic burden, while its influence on macrophage lipid accumulation remains more limited.
- The parameters *ν, θ*, and *λ* exhibit generally weak influence across most model outputs, indicating that they do not play a dominant role in shaping the system dynamics.
- The parameter *κ* exhibits its strongest influence on the macrophage populations *M* and *A*_*M*_; however, its effect remains substantially weaker compared to the dominant influence of the proliferation rate *ρ* and the emigration rate *γ*.
- These results are consistent with the asymptotic fast-slow structure established in Section 4. In particular, the proliferation rate *ρ* and the emigration rate *γ* emerge as the dominant parameters influencing the slow macrophage variables *M* and *A*_*M*_, whereas the efferocytosis rate *η* exerts its strongest influence on the apoptotic variables *P* and *A*_*P*_.

Overall, the PRCC analysis reveals that plaque dynamics are governed by a structured hierarchy of biological mechanisms. Macrophage proliferation (*ρ*) acts as the dominant driver of system behaviour, emigration (*γ*) provides strong suppression, and efferocytosis (*η*) introduces selective regulation of apoptotic cell clearance. This hierarchy has direct biological implications: interventions targeting proliferation are expected to produce broader system-wide effects, whereas modulation of efferocytosis is likely to yield more selective changes in apoptotic burden, with comparatively weaker effects on the remaining model outputs.

The PRCC analysis identifies *ρ* and *γ* as the dominant parameters governing macrophage accumulation and live macrophage lipid content, while *η* has its most pronounced influence on the apoptotic variables *P* and *A*_*P*_. Notably, the dominance of *ρ* is fully consistent with the bifurcation result established in Section 4, where the proliferation rate determines the critical threshold *ρ*_*c*_ = 1+*γ* for the transcritical bifurcation at infinity. The following subsections use these dominant parameters to illustrate how the bifurcation structure manifests across the parameter space, connecting the analytical result to concrete biological outcomes.

### 5.2 Parameter sweeps near the bifurcation threshold

The most efficient parameters (*ρ, γ*, and *η*) identified by the PRCC analysis are now examined through targeted simulations centred on the neighbourhood of *ρ* = *ρ*_*c*_. The aim is to show concretely how the analytical transition identified in Section 4 is reflected in the dynamic and steady state behaviour of the system across these parameter ranges.

#### 5.2.1 Threshold dynamics and blow-up behaviour induced by *ρ*

To examine how the transcritical bifurcation is obvious dynamically, we analyse the time evolution of the key state variables *M* (*t*), *P* (*t*), *A*_*M*_ (*t*), *A*_*P*_ (*t*), and *N* (*t*) for varying values of *ρ* near and beyond the critical threshold *ρ* = *ρ*_*c*_.

When *ρ* remains below the critical threshold, all variables approach biologically meaningful steady states, consistent with the stability analysis of Section 3.1 (see Figure 5). To further illustrate the dynamics near and at the bifurcation threshold, Figure 6 presents the time evolution for *ρ* = 1.14 and *ρ* = *ρ*_*c*_ = 1.15. For *ρ* = 1.14, all variables converge to a finite steady state, while at *ρ* = *ρ*_*c*_ = 1.15 the finite equilibrium is lost and the variables grow without bound, consistent with the transcritical bifurcation at infinity established analytically in Section 4.

**Fig. 5.**
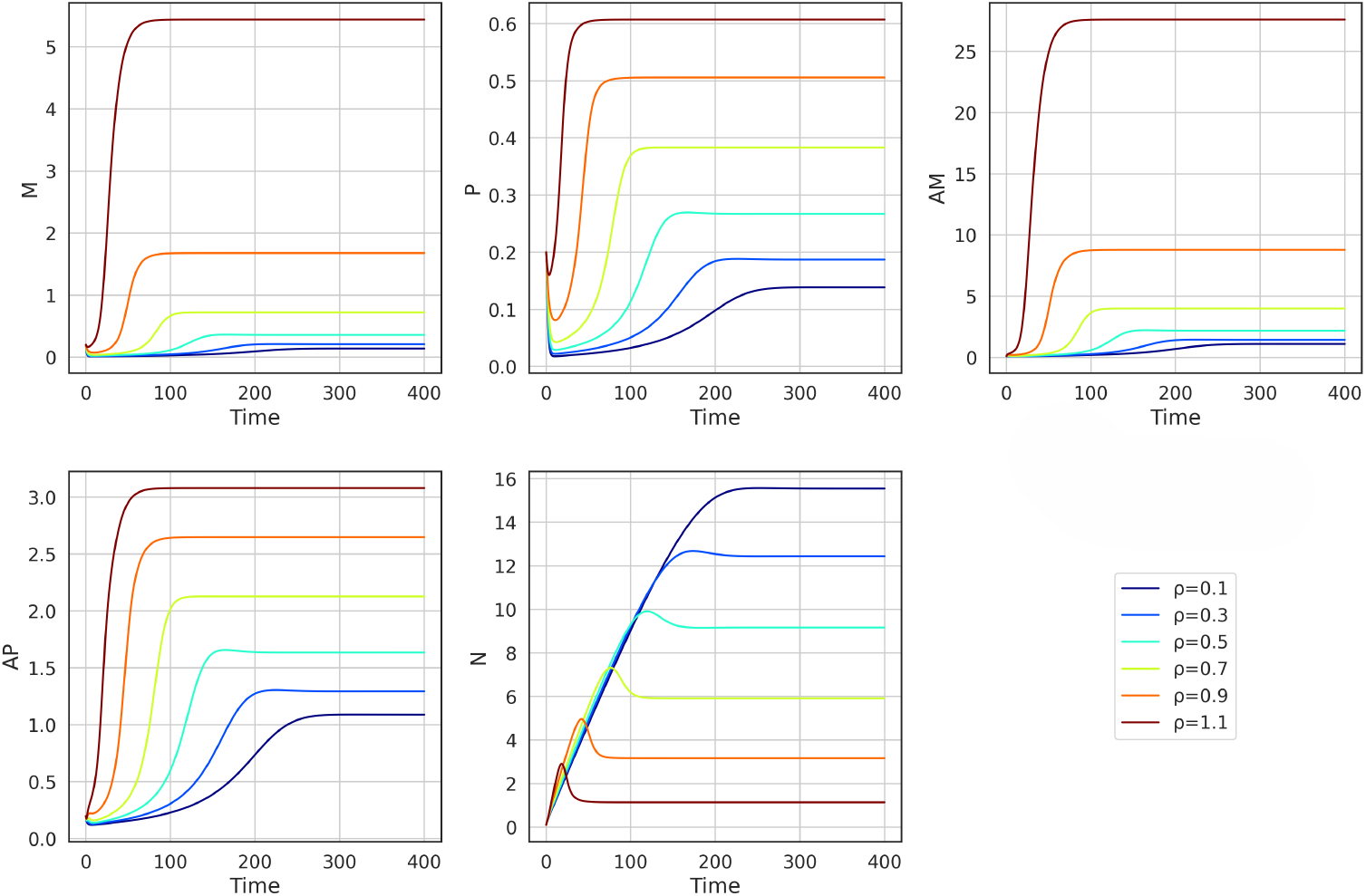
Time evolution of the system variables *M*, *P*, *A*_*M*_, *A*_*P*_, and *N* for varying values of the parameter *ρ*, from 0.1 to 1.1 with a 0.2 step-difference, all strictly below the bifurcation threshold *ρ*_*c*_ = 1.15. As *ρ* increases toward *ρ*_*c*_, the system converges to progressively larger steady state values with slower convergence

**Fig. 6.**
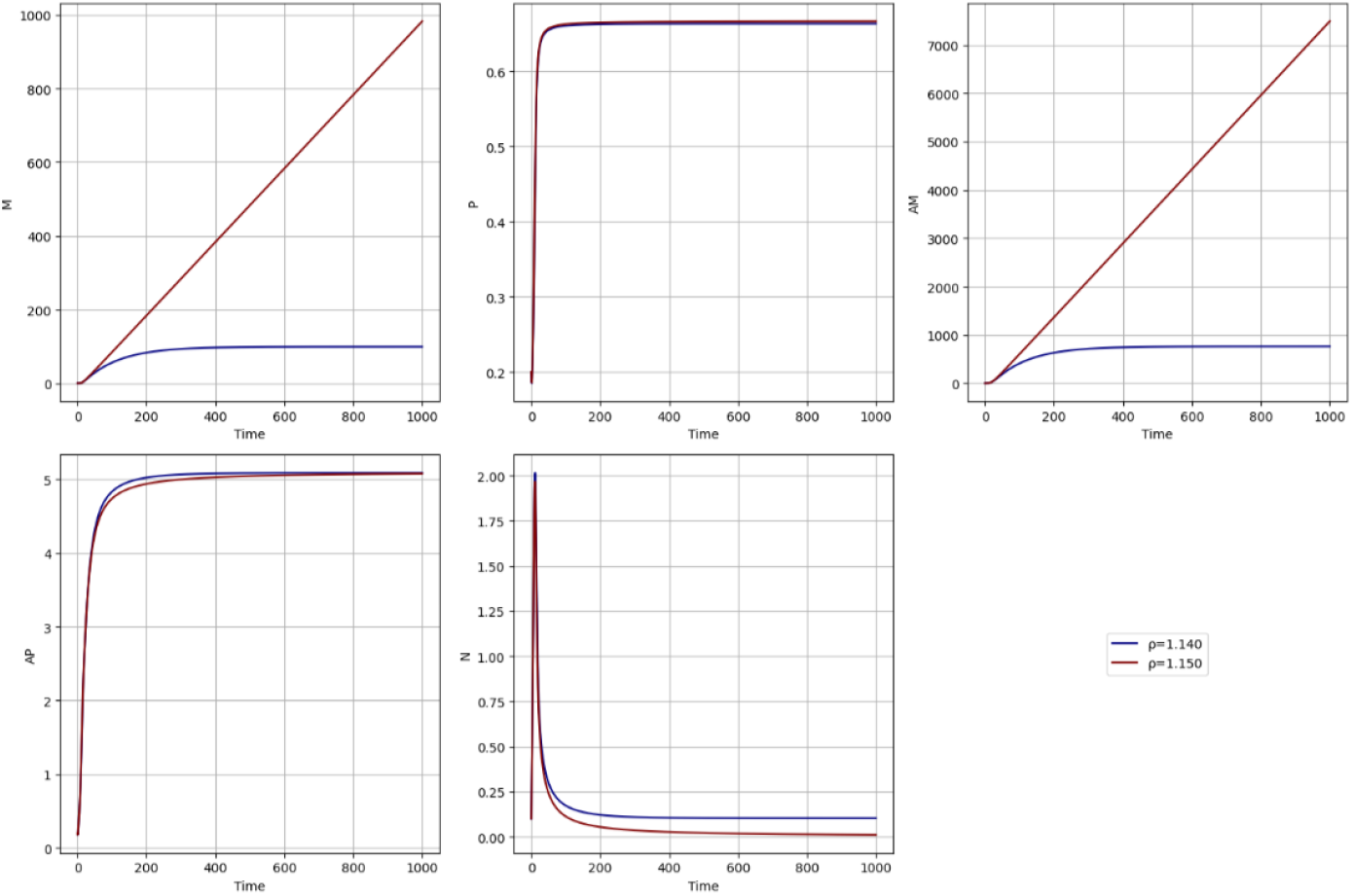
Time evolution of the system variables *M*, *P*, *A*_*M*_, *A*_*P*_, and *N* for *ρ* = 1.14 (below threshold) and *ρ* = *ρ*_*c*_ = 1.15 (at threshold). The simulations demonstrate a qualitative shift in behaviour at the bifurcation threshold. For *ρ* = 1.14, all variables converge to a finite steady state, whereas at *ρ* = *ρ*_*c*_ = 1.15 the variables grow without bound, consistent with the transcritical bifurcation at infinity established analytically in Section 4. Results for *ρ* > *ρ*_*c*_ are not shown due to the extreme scale of the unbounded growth, which would obscure the dynamics below the threshold

From a biological perspective, such runaway dynamics reflect a failure of immune regulation in which macrophage proliferation overwhelms the combined effects of apoptosis and emigration. Within the asymptotic regime analysed here, this corresponds to a state of chronic unresolved inflammation, consistent with the progressive plaque destabilisation observed in advanced atherosclerosis. Importantly, the transcritical structure of this transition suggests that restoring the proliferation-emigration balance below *ρ*_*c*_ may promote a return toward regulated macrophage dynamics by reassigning local stability to the finite regulatory state.

#### 5.2.2 Regulatory influence of *η* and *γ* on plaque macrophage dynamics

We illustrate the time evolution of model (1) variables *M*, *P*, *A*_*M*_, *A*_*P*_, and *N* for varying values of the parameter *η*, determining how efficiently live macrophages clear apoptotic cells and their associated lipid content (*A*_*P*_) from the live macrophage lipid content pool (*A*_*M*_) in Figure 7. All other parameters, except *η*, are fixed as in Table 1, and *ρ* fixed at 0.8, well within the stable regime *ρ* < *ρ*_*c*_ = 1.15. The specific values of *η* used in the simulations correspond to those listed in the accompanying table.

**Fig. 7.**
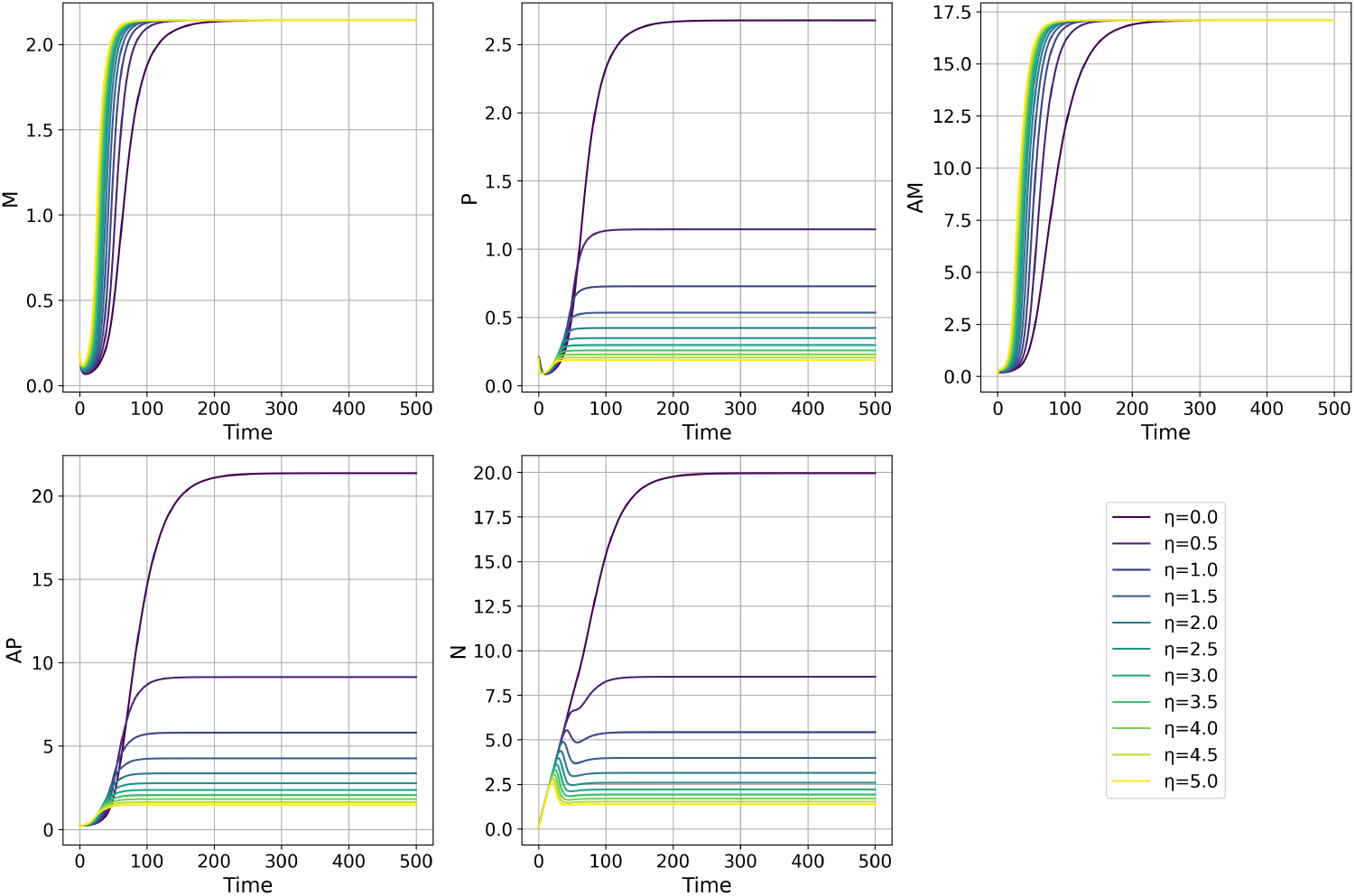
Time evolution of the system variables *M*, *P*, *A*_*M*_, *A*_*P*_, and *N* for varying values of the parameter *η*

As *η* increases, a marked decrease in the steady state values of *P, A*_*P*_, and *N* is observed, while *M* and *A*_*M*_ remain largely unaffected across the range of *η*. The fast-slow analysis of Section 4 provides a natural explanation for this observation: the fast variables *P*, *A*_*P*_, and *N* rapidly approach *η*-dependent quasi-steady states, whereas the reduced slow subsystem governing *M* and *A*_*M*_ is, to leading order, independent of *η*. Biologically, this indicates that while efferocytosis effectively reduces apoptotic burden and necrotic core lipid content, macrophage accumulation is primarily regulated by the proliferation and emigration balance characterised by *ρ*_*c*_. A low *η* results in elevated levels of *A*_*P*_ and *N*, potentially reflecting an unresolved inflammatory state, whereas high *η* facilitates faster resolution of apoptotic burden while leaving macrophage dynamics unchanged.

Figure 8 depicts the time evolution of the system variables *M*, *P*, *A*_*M*_, *A*_*P*_, and *N* for varying values of the macrophage emigration rate *γ*. The parameter *ρ* is fixed at 0.8, while all other parameters are held constant as listed in Table 1. As *γ* increases, a general decline in the steady state values of *M*, *P*, *A*_*M*_, and *A*_*P*_ is observed, reflecting the enhanced emigration and reduction of macrophage populations involved in pathogen recognition and immune activation. Interestingly, as *γ* rises, *N* tends to increase, consistent with a compensatory buildup of downstream factors. It is because the drop in *M* weakens *M*-mediated clearance (−*θMN*), slowing the decline of *N* and, when *νA*_*P*_ > *θMN*, allowing it to grow. These results highlight the critical role of *γ* in modulating the balance between macrophage accumulation and clearance, thereby shaping the overall dynamics of the immune response.

**Fig. 8.**
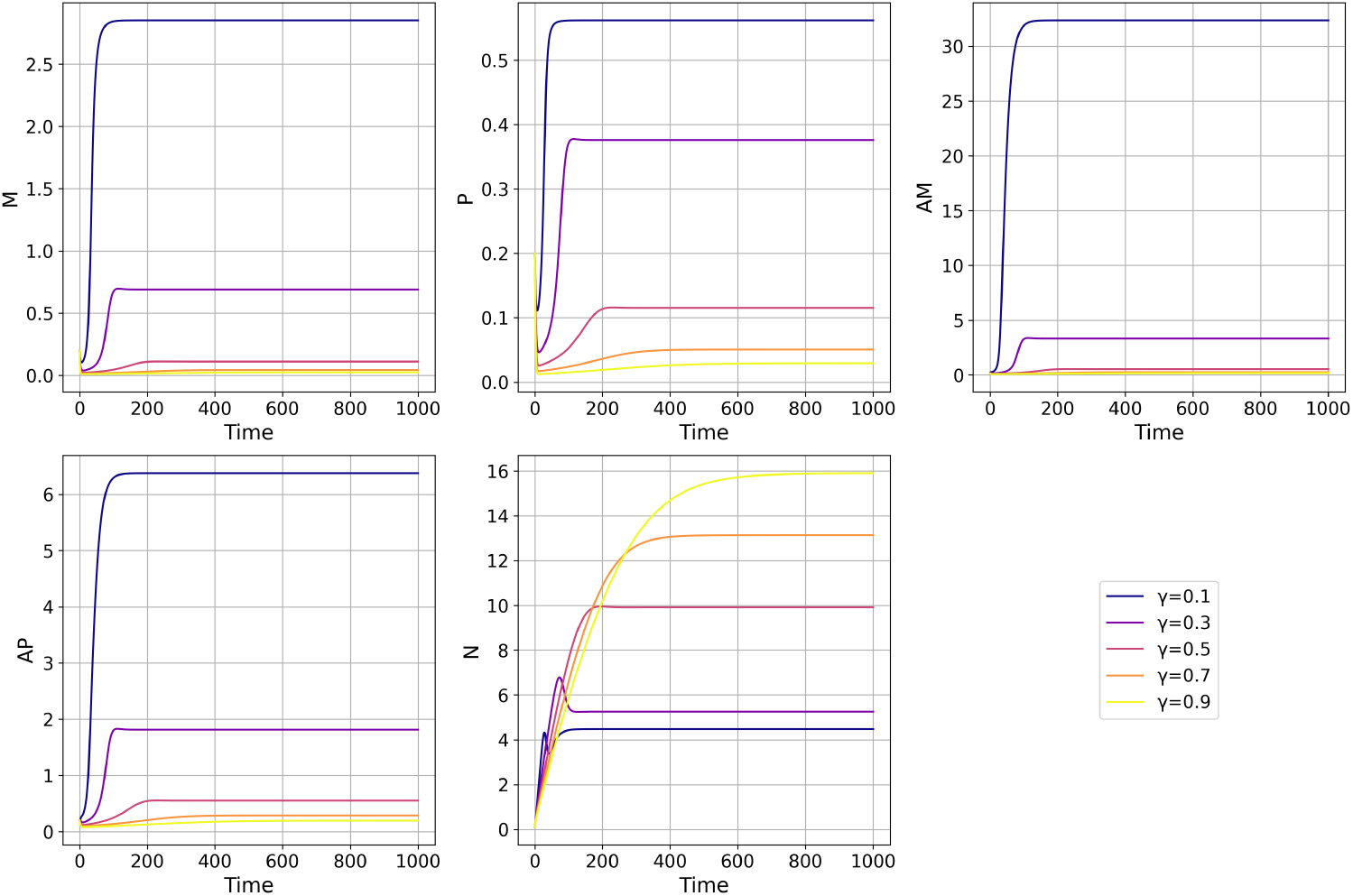
Time evolution of the system variables *M*, *P*, *A*_*M*_, *A*_*P*_, and *N* for varying values of the macrophage emigration rate *γ*

#### 5.2.3 Steady state heatmap analysis

The steady state behaviour of the system is examined through heatmaps of the macrophage population *M*\* and necrotic core lipid content *N*\* across the (*ρ, η*) parameter space, and the live macrophage lipid content 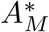 and necrotic core lipid content *N*\* across the (*ρ, γ*) parameter space.

Figure 9 presents heatmaps of the equilibrium macrophage population *M*\* and necrotic core lipid content *N*\* as functions of the proliferation rate *ρ* and efferocytosis rate *η*, both identified by the PRCC analysis as dominant regulators of system behaviour. *M*\* remains low across most of the (*ρ, η*) parameter space, with a sharp increase near the critical threshold *ρ* = *ρ*_*c*_, independent of *η*, confirming that macrophage accumulation is primarily driven by *ρ*. In contrast, *N*\* increases with *ρ* and decreases with *η*, highlighting that effective efferocytosis is necessary to prevent necrotic core expansion regardless of the proliferation rate.

**Fig. 9.**
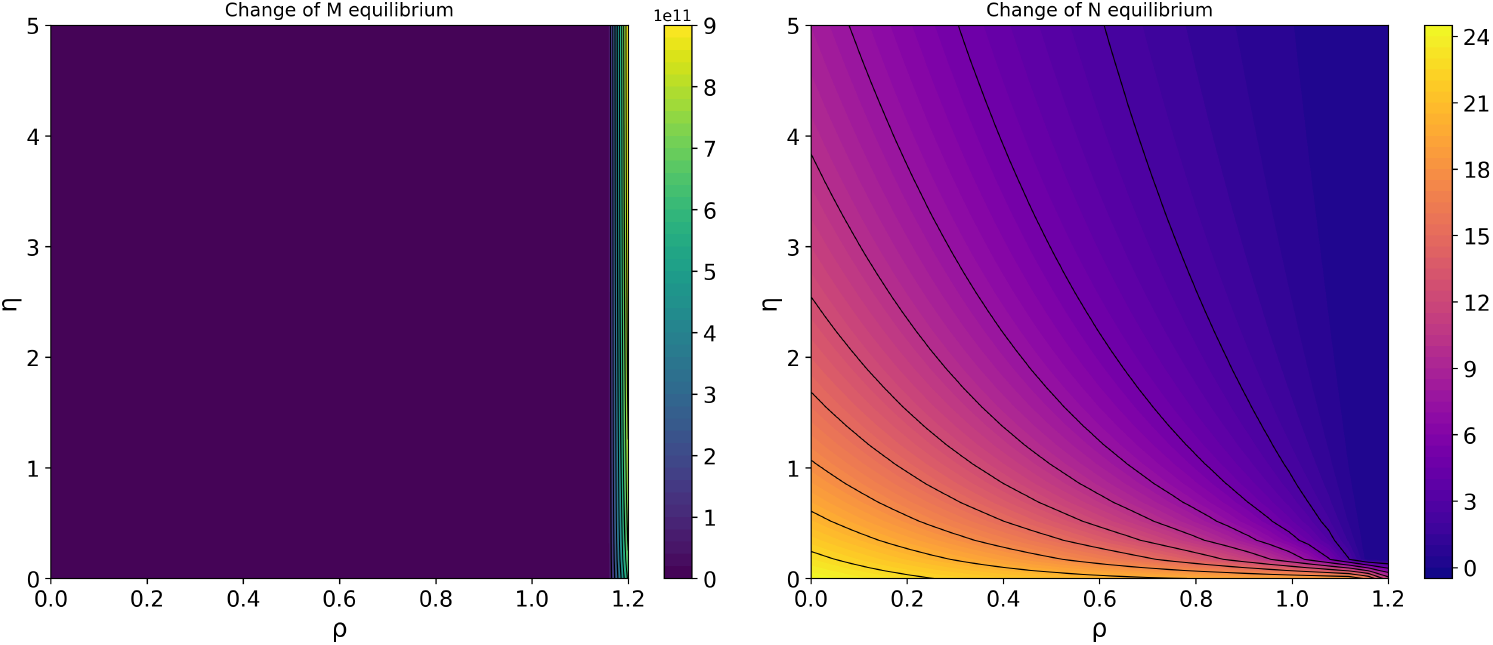
Heatmap showing the equilibrium values of variables *M* and *N* as functions of the parameters *ρ* and *η*. In the (*ρ, η*) parameter space, macrophage *M* remains low and essentially insensitive to *η* until a sharp increase at the critical threshold *ρ*_*c*_ = 1.15, while the necrotic core *N* rises with *ρ* and falls with *η*

Figure 10 presents heatmaps of the live macrophage lipid content 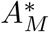 and necrotic core lipid content *N*\* as functions of the proliferation rate *ρ* and emigration rate *γ*. 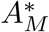 remains low across most of the (*ρ, γ*) parameter space, with a steep increase along the threshold line *ρ* = *ρ*_*c*_, confirming that a higher emigration rate requires a higher proliferation rate to trigger unbounded macrophage accumulation, consistent with the transcritical bifurcation established in Section 4. In contrast, *N*\* exhibits a non-monotone response to *γ*: at low *γ, N*\* increases substantially with *ρ*, but sufficiently high *γ* suppresses *N*\* regardless of *ρ*. This is because emigrating macrophages carry lipid out of the plaque before it can be deposited into the necrotic core; at high emigration rates, this removal mechanism dominates, suppressing necrotic core lipid content even under high proliferation rates. This reveals a regulatory decoupling between macrophage accumulation and necrotic core growth, where emigration can suppress necrotic core lipid content even under high proliferation rates.

**Fig. 10.**
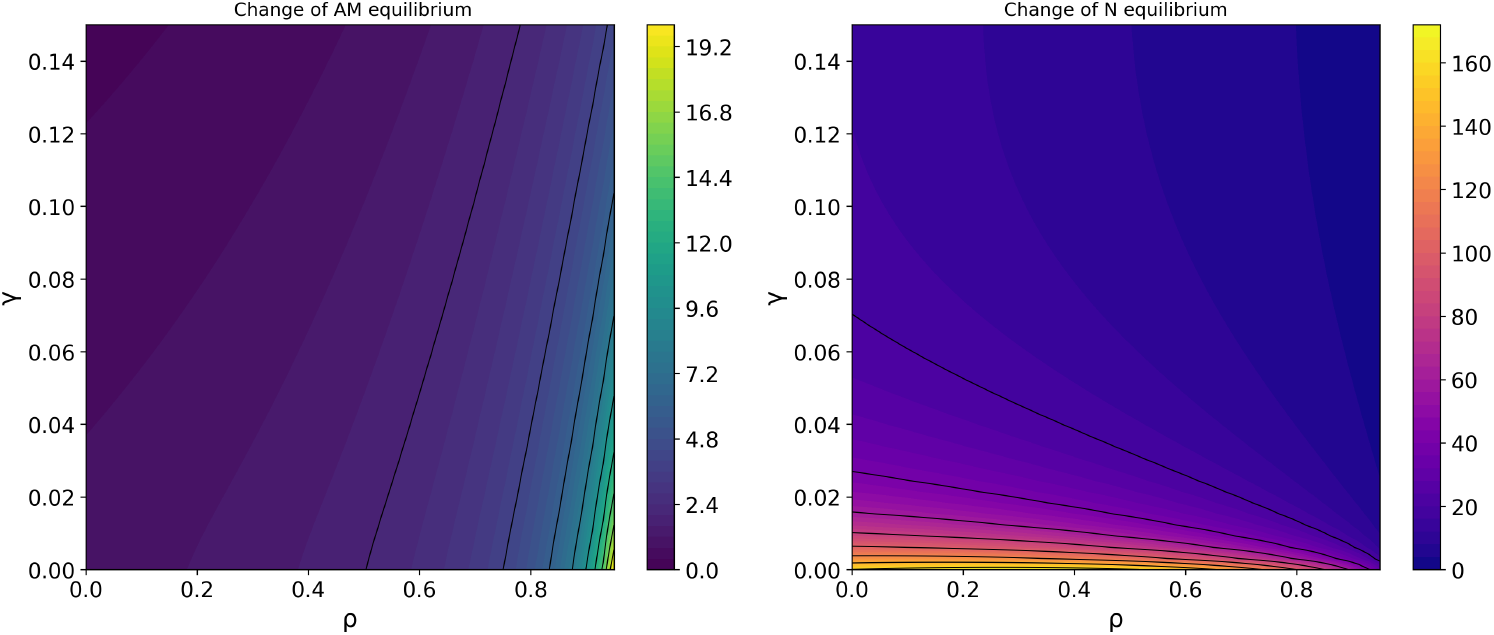
Heatmap showing the equilibrium values of variables *A*_*M*_ and *N* as functions of the parameters *ρ* and *γ*

## 6 Discussion

The role of macrophage proliferation in atherosclerotic plaque progression remains paradoxical. While some studies suggest that proliferation contributes to plaque instability Rosenfeld (2014); Wang et al. (2017), others propose that proliferative macrophage subpopulations may be protective Yurdagul (2022); Robbins et al. (2013). For the monocyte-derived macrophage population, Robbins et al. (2013) demonstrated that local proliferation accounts for approximately 87% of newly accumulated lesional macrophages in established atherosclerosis, thereby identifying proliferation as the dominant mechanism of macrophage accumulation. Our analysis of the asymptotic large-macrophage regime provides a mechanistic resolution of this paradox: proliferation is neither inherently pathological nor protective, but its effect depends entirely on the balance with macrophage emigration. When this balance tips beyond the critical threshold *ρ*_*c*_ = 1 + *γ* established in Section 3, the finite regulatory state loses its stabilising influence, and the runaway accumulation regime becomes the dominant dynamical tendency. This is consistent with the broader view of Libby et al. (2009) that targeting specific macrophage-driven processes may be atheroprotective, and provides a precise mathematical criterion for identifying when such targeting is most needed.

In this study, we revisited the macrophage proliferation model of Chambers et al. (2024b), which describes the interplay between lipid accumulation and macrophage dynamics in atherosclerotic plaque development. While Chambers et al. restricted their analysis to *ρ* < *ρ*_*c*_, regarding the regime *ρ ≥ ρ*_*c*_ as unphysical, our bifurcation analysis shows that the system remains mathematically well-structured in this regime. In particular, we identify a *transcritical bifurcation at infinity* at *ρ* = *ρ*_*c*_, providing a rigorous dynamical systems characterisation of this transition that goes beyond the steady state description of Chambers et al. (2024b). Our contribution is thus twofold: (i) we extend the mathematical analysis of the model into a previously unexplored parameter regime, and (ii) we identify the precise bifurcation mechanism governing the loss of plaque stability, offering new insight into the conditions under which plaque growth becomes uncontrolled and, crucially, the conditions under which local stability of the finite regulatory state may be restored. Together, these findings identify a previously unrecognised dynamical threshold that offers a new perspective on macrophage-driven plaque progression, opening a pathway for future experimental investigation of whether biological systems operate near such critical balances.

While standard local bifurcation techniques (see Kuznetsov (2004)) are well-suited for finite equilibria, the characterisation of transcritical bifurcations at infinity requires more advanced global methods. In particular, the fast-slow decomposition of the system, combined with centre manifold theory, allows us to track the system behaviour as the macrophage population grows without bound. Unlike classical transcritical bifurcations occurring at finite equilibria, where two equilibrium branches intersect and exchange stability Kuznetsov (2004), a transcritical bifurcation at infinity occurs through an exchange of stability between the finite positive equilibrium branch and the equilibrium at infinity. This exchange indicates a potentially reversible transition within the model: bringing the proliferation-emigration balance below *ρ*_*c*_ restores local stability to the finite regulatory state. As shown in Figures 1 and 2, the Jacobian determinant changes sign and one eigenvalue approaches zero precisely at *ρ* = *ρ*_*c*_, providing numerical evidence for this transition. The analytical proof of this result is established in Section 4.

Instability arises when the proliferation rate exceeds the combined rate of apoptosis and emigration, that is, when *ρ* > *ρ*_*c*_. Below this threshold, the system maintains a stable regulatory equilibrium; above it, the *transcritical bifurcation at infinity* marks the transition to a regime in which the macrophage population and lipid content grow without bound while *ρ* remains above the critical threshold, as illustrated in Figures 5 and 6. The sensitivity analysis further reveals that while the efferocytosis rate *η* significantly influences plaque severity, it does not alter the bifurcation threshold *ρ*_*c*_. This can be understood from the system’s fast-slow structure: efferocytosis acts only through the fast variables, whereas the bifurcation is governed by the slow macrophage dynamics. We emphasise that this fast-slow separation is a feature of the asymptotic regime *M* → ∞, *u* → 0^+^ analysed here; outside this limit, the apoptotic and necrotic dynamics are fast only if the corresponding lipid uptake rates are taken sufficiently large, and the model does not otherwise exhibit this timescale separation. Consequently, while efferocytosis modulates necrotic core growth within the stable regime (see Figure 7), it cannot prevent the fundamental transition when *ρ* > *ρ*_*c*_.

These results suggest potential clinical implications within the asymptotic regime analysed here. The characterisation of *ρ* = *ρ*_*c*_ as a transcritical bifurcation at infinity reveals not only that solutions grow without bound for *ρ* > *ρ*_*c*_, but precisely why this transition occurs and what governs it. As established in Section 4, the bifurcation is driven by the slow macrophage dynamics, identifying macrophage proliferation and emigration as the primary determinants of long-term plaque fate in this asymptotic regime. Importantly, the transcritical structure of this transition suggests that restoring the proliferation-emigration balance below *ρ*_*c*_ may promote a return toward regulated macrophage dynamics by restoring local stability to the finite regulatory state. Therapeutic strategies targeting both proliferation and emigration may therefore offer a more robust approach than targeting efferocytosis alone, provided the system operates within or near the asymptotic regime considered here. Furthermore, since necrotic core size is clinically measurable, its relationship with the proliferation-emigration balance may provide a basis for future investigation of patient-specific proximity to *ρ*_*c*_ and the associated risk of plaque destabilisation.

Recent spatially resolved and dual-structured models by Chambers and co-workers have significantly advanced our mechanistic understanding of early atherosclerotic lesions, particularly the interplay between LDL retention, HDL capacity, lipid loading, and phenotype transitions within macrophage populations Chambers et al. (2024a, 2025). These models provide a detailed description of micro-scale behaviour in early-stage plaques and consistently predict bounded steady states in the absence of macrophage proliferation. In contrast, the model we analyse operates at a more aggregated scale in which macrophage gain and loss processes are encoded through effective proliferation and emigration rates based on Section 3.2.3. Our bifurcation analysis shows that this reduced system exhibits a *transcritical bifurcation at infinity* in the asymptotic large-macrophage regime, identifying a mathematically sharp threshold beyond which plaque growth becomes unbounded as in Figure 3. Hence, our results complement the detailed early-lesion models of Chambers et al. (2024a, 2025) by revealing how proliferation-driven mid-stage dynamics may lead to catastrophic plaque expansion if the proliferation-emigration balance is not restored, consistent with the onset of myocardial infarction.

Several natural extensions of this work could be explored in future. First, linear macrophage proliferation is considered in the present model, which describes the fundamental growth-emigration balance underlying the *transcritical bifurcation at infinity*. Biological evidence suggests, however, that macrophage replication may be regulated by density-dependent mechanisms, including contact inhibition and resource limitation Vaughan-Jackson et al. (2022). Incorporating a logistic-type proliferation term would naturally bound the unbounded growth regime and may enrich the bifurcation structure identified here. Rather than eliminating the underlying proliferation-emigration balance, such an extension would be expected to transform the unbounded growth regime into a saturated but still dynamically distinct state, providing a more physiologically realistic framework for predicting plaque fate. Second, the present model describes an aggregate population of monocyte-derived macrophages. However, Shankman et al. (2015) demonstrated that smooth muscle cell-derived macrophage-like cells constitute a significant fraction of lesional cells and exhibit distinct proliferative and apoptotic dynamics. Extending the lipid-structured framework to incorporate multiple macrophage subpopulations with different kinetic properties would provide a more complete description of plaque cellular composition and may reveal a richer bifurcation structure. Finally, establishing quantitative links between model parameters and clinically measurable quantities such as necrotic core size would enhance the translational value of this framework for identifying patients at high risk of plaque destabilisation.

## Appendix A Main model formulation

The original PIDE model given in Chambers et al. (2024b) is

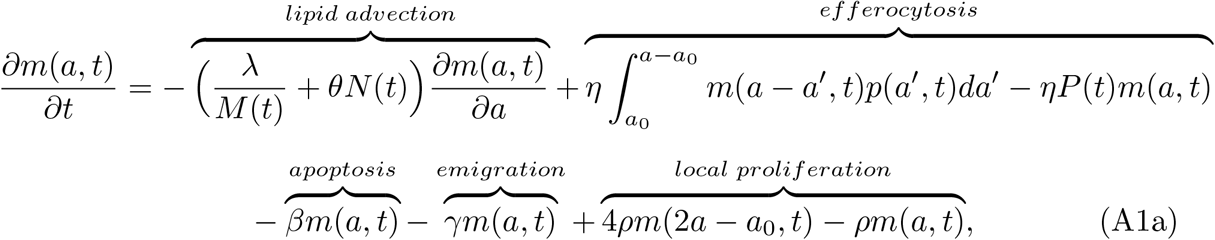

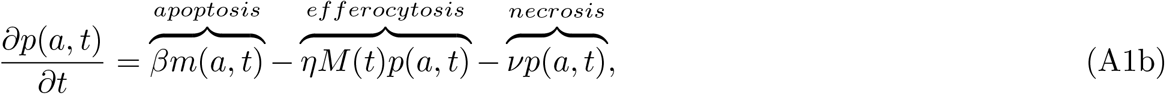

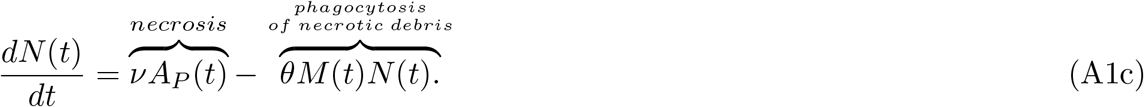

Here, the density numbers of live and apoptotic macrophages in the lesion, denoted by *m*(*a, t*) and *p*(*a, t*), are functions of lipid content *a* ≥ a_0_ and time *t* ≥ 0. The constant *a*_0_ corresponds to the baseline lipid content present in the membranes of macrophage cells, and the integral in (A1a) is defined as zero for *a* < 2*a*_0_. The integral term in (A1a) represents lipid uptake through efferocytosis rather than macrophage proliferation. The lower integration limit is therefore *a*_0_, since apoptotic macrophages contain at least the baseline intracellular lipid content *a*_0_, and only this lipid is transferred to the engulfing macrophage during efferocytosis. Additionally, the necrotic core’s lipid content is given by *N* (*t*). To determine the total populations of live and apoptotic macrophages, denoted *M* (*t*) and *P* (*t*), the densities *m*(*a, t*) and *p*(*a, t*) are integrated over all possible lipid contents:

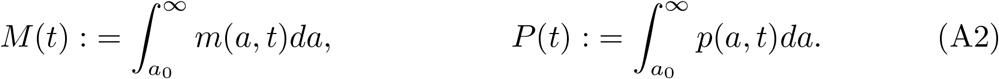

Similarly, the total lipid quantities carried by the live and apoptotic macrophage populations, denoted *A*_*M*_ (*t*) and *A*_*P*_ (*t*), are obtained by integrating the lipid-weighted densities:

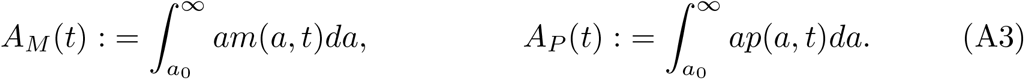

The remaining parameters are positive constants that govern the system dynamics: lipid uptake rates include *λ* (the net rate of lipid uptake via LDL/HDL), *θ* (the rate of necrotic lipid uptake), and *η* (the rate of apoptotic lipid uptake). Macrophage dynamics are regulated by *β* (apoptosis rate), *γ* (emigration rate), *ρ* (proliferation rate), and *ν* (secondary necrosis rate). For a comprehensive derivation and biological interpretation of these terms, including their roles in lesion progression, we direct the reader to Chambers et al. (2024b).

### A.1 Boundary conditions

The model (A1) is supplemented by a boundary condition at the endogenous lipid level, *a* = *a*_0_, that describes the recruitment of new macrophages from the bloodstream. This is given by

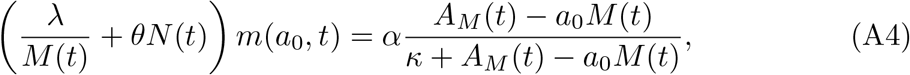

where *t* > 0. The parameter *κ* represents the lipid threshold for half-maximal recruitment in live cells, while *α* denotes the maximum recruitment rate.

### A.2 Initial conditions

The initial conditions follow Chambers et al. (2024b), with lipid distributions given by half-normal functions centred at the endogenous lipid level *a*_0_:

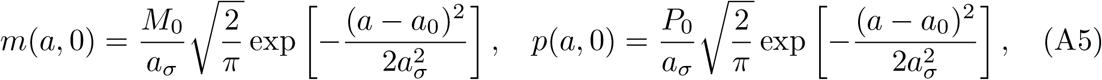

where *a*_*σ*_ *>* 0 is a constant that determines the spread of the initial lipid load. Here, the lesion was initialised with no necrotic core (*N* (0) = 0), meaning that all macrophages initially contain only endogenous lipid. Furthermore, the initial number of apoptotic macrophages (*P* (0) = *P*_0_) is set to be smaller than the live population (*M* (0) = *M*_0_), satisfying *M*_0_ > *P*_0_ via the relation 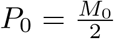.

The initial macrophage population, *M*_0_, and lipid moments will be specified in Section A.3, along with the ODE initial conditions, to ensure consistency across the model formulation.

### A.3 ODE reduction

The non-local PIDE system (A1) shows a comprehensive description of lipid distribution within the macrophage population. However, to better understand the overall dynamics of plaque formation and simplify the analysis, a reduced system of ordinary differential equations (ODEs) is given by integrating the PDEs in (A1) with respect to the lipid variable *a* ∈ (*a*_0_, ∞).

If we first integrate (A1a) over all lipid contents,

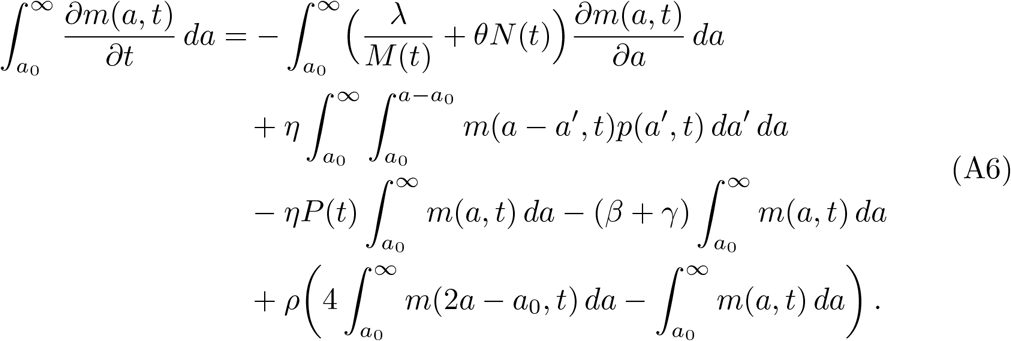

For (A6), the key simplifications are gotten as

- Advection term: 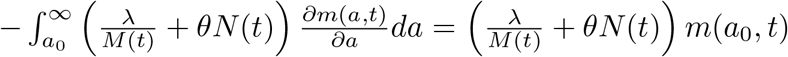 (leads to the boundary condition)
- Efferocytosis non-local part: 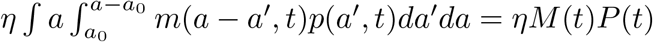
- Proliferation lipid synthesis: *ρM* (*t*).

Using the same integration approach for (A1b) and gathering the other ODE equations, we yield

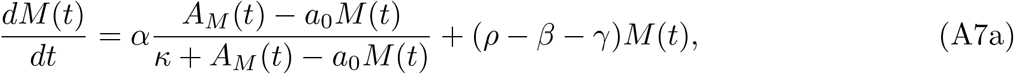

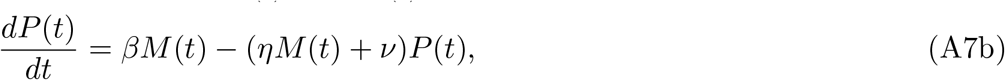

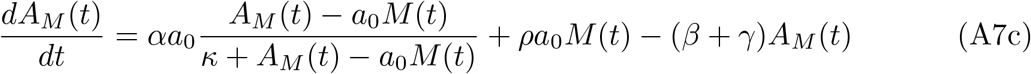

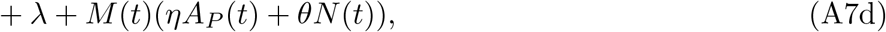

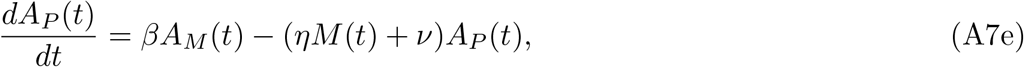

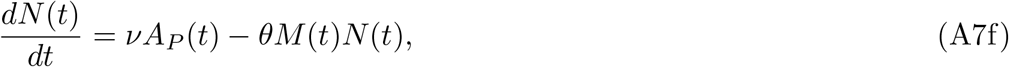

where *A*_*M*_ (*t*) and *A*_*P*_ (*t*) can be defined by integration *aM* (*t*) and *aP* (*t*) respectively. The initial conditions for the ODE subsystem (A7) are determined by integrating the initial lipid distributions. This ensures mathematical consistency between the integro-PDE and ODE formulations. Specifically,

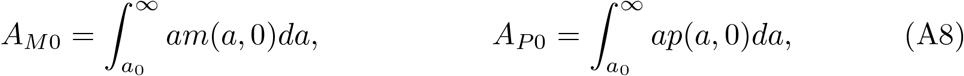

*m*(*a*, 0) and *p*(*a*, 0) are given by the half-normal distributions in (A5). For these distributions, the integrals evaluate to

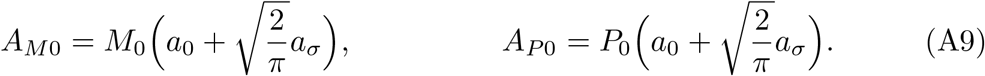

Here *M*<u>_0_ i</u>s derived using the boundary condition given in (A4), and defined as 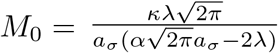. For biologically meaningful, 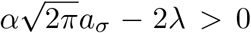 must be satisfied to ensure *M*_0_ > 0.

#### A.4 Nondimensionalisation

To simplify analysis and reduce the number of free parameters, here we give the rescaled version of the model equations (A7). The rescaled macrophage lipid content is defined as:

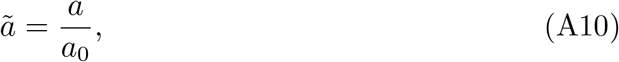

where *a* is the total lipid content and *a*_0_ is the basal lipid content. And time by the mean macrophage lifespan *β*^−1^ as

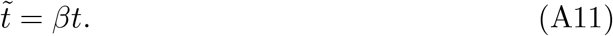

Based on these descriptions, the other variables of (A7) are scaled as

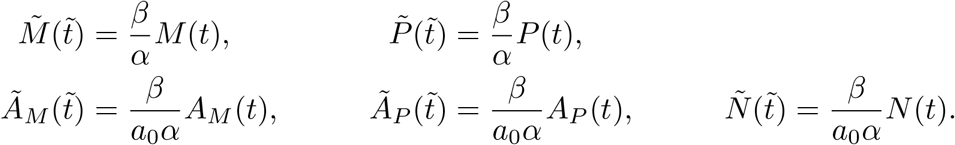

The dimensionless parameters are defined as:

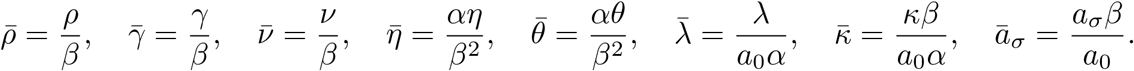

## Appendix B Technical details

Additional technical details of the centre manifold reduction, including justification of smoothness and higher-order expansions, can be found in this appendix.

### Theorem 4

(Fenichel’s Invariant Manifold Theorem Fenichel (1979); Jones (1995)) *Consider the singularly perturbed system*

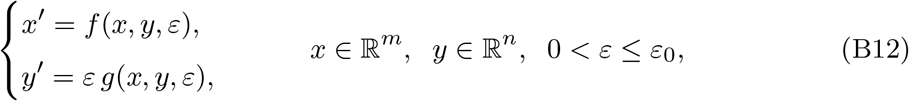

*where f and g are C*^*r*^ (*r* ≥ 2) *vector fields defined on an open set U* ⊂ ℝ ^*m*+*n*^ *×* [0, *ε*_0_]. *Define the* critical manifold

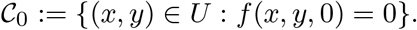

*Assume that:*

1. *C*_0_ *is a compact C*^*r*^ *manifold (possibly with boundary);*
2. *for every* (*x, y*) ∈ *C*_0_, *the Jacobian D*_*x*_*f* (*x, y*, 0) *has no eigenvalues with zero real part, so that C*_0_ *is* normally hyperbolic;
3. *the hyperbolicity is* uniform *on C*_0_, *i*.*e. there exists α* > 0 *such that all eigenvalues λ of D*_*x*_*f* (*x, y*, 0) *satisfy* |ℜλ| ≥ *α for all* (*x, y*) ∈ *C*_0_.

*Then there exists ε*_1_ > 0 *such that, for each ε* ∈ (0, *ε*_1_], *the full system* (B12) *admits a locally invariant manifold C*_*ε*_ *with the following properties:*

1. *C*_*ε*_ *is a C*^*r*^ *manifold diffeomorphic to C*_0_, *and C*_*ε*_ *lies within O*(*ε*) *of C*_0_ *in the C*^0^ *topology;*
2. *the flow on C*_*ε*_ *is C*^*r*^*-conjugate to the reduced (slow) flow on C*_0_, *obtained by substituting x* = *h*_0_(*y*) *into y*′ = *εg*(*x, y*, 0);
3. *if all eigenvalues of D*_*x*_*f* (*x, y*, 0) *have negative real part (i*.*e. C*_0_ *is attracting), then C*_*ε*_ *is also an attracting invariant manifold;*
4. *the stable and unstable fibers of C*_*ε*_ *depend smoothly on ε and form C*^*r*^ *foliations of a neighborhood of C*_0_.

## Appendix C Detailed eigenvalue behaviour for small values of *ρ*

To provide a clearer view of the stability behaviour near the initial parameter range, Figure C1 shows the eigenvalues over a restricted interval of *ρ*. In the corresponding figure in the main text, some of the real parts decrease rapidly to large negative values as *ρ* increases, which substantially enlarges the vertical scale and makes the sign of the eigenvalues near the beginning of the parameter interval difficult to distinguish visually.

The enlarged view confirms that the real parts of all eigenvalues remain strictly negative throughout the displayed range. Thus, the apparent proximity of some eigen-value curves to the zero axis in the full-range plot is a consequence of the plotting scale rather than a loss of stability.

**Fig. C1.**
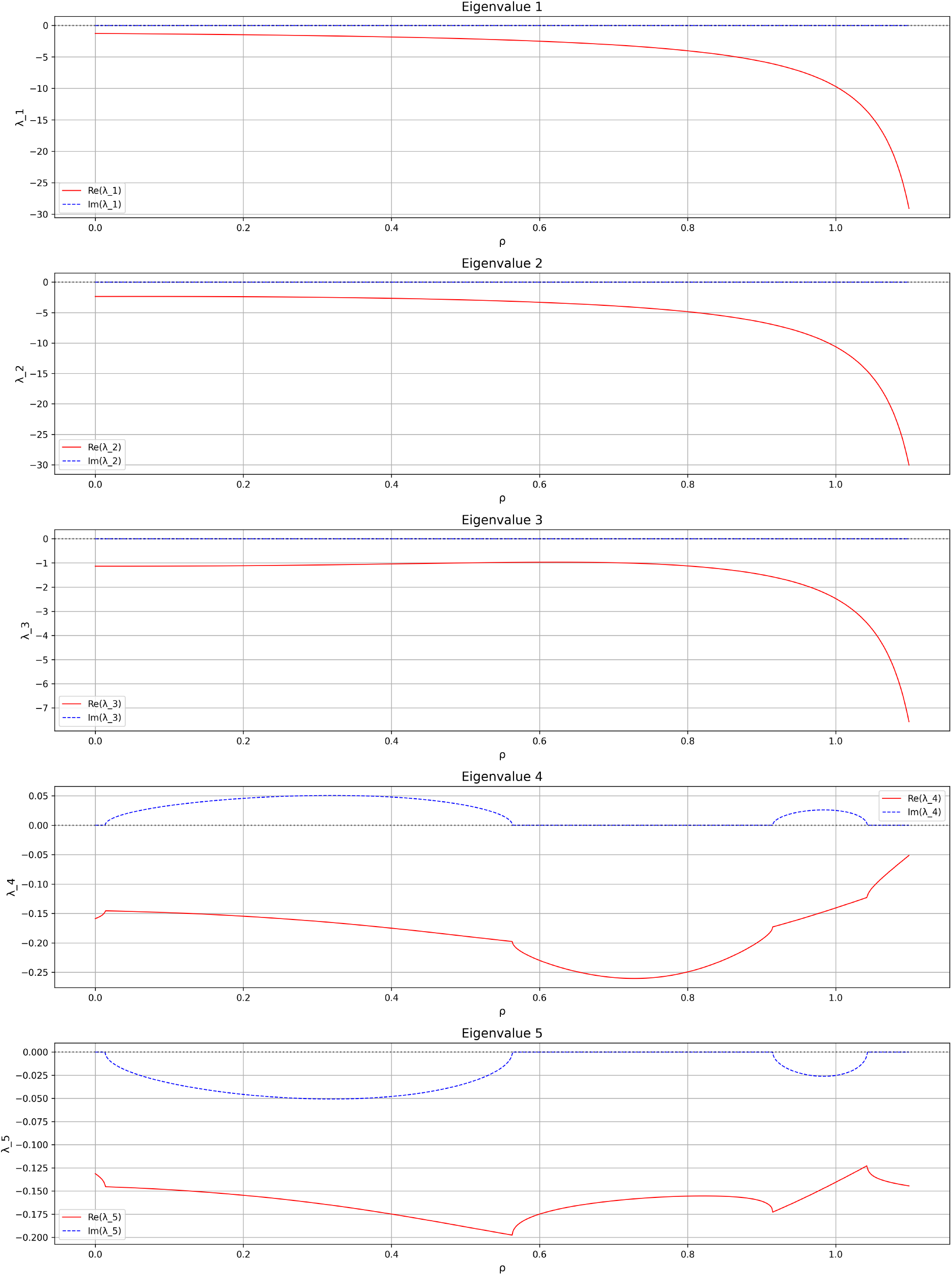
Detailed eigenvalue behaviour over a restricted range of *ρ*. The solid red curves represent the real parts of the eigenvalues, while the dashed blue curves represent their imaginary parts. The enlarged scale confirms that all real parts remain negative over this interval. This restricted-range representation complements the full-range eigenvalue plot in the main text, where the rapid decrease of some eigenvalues to large negative values compresses the curves near the zero axis.

## Acknowledgments

We would like to express our sincere appreciation to Assoc. Prof. Dr Adem Atıcı for his insightful comments regarding cardiovascular disease. We also thank Prof. Jonathan A. Sherratt for his insightful feedback and constructive suggestions.

## Declarations

### Data Availability

No underlying data are available for this article, since no datasets were generated or analysed during this study.

